# Identification of a mesodermal progenitor for the pro-definitive angio-hematopoietic lineage

**DOI:** 10.1101/2024.08.24.609533

**Authors:** Tomi Lazarov, Pierre-Louis Loyher, Hairu Yang, Zi-Ning Choo, Zihou Deng, Sonja Nowotschin, Ying-Yi Kuo, Ting Zhou, Araitz Alberdi-Gonzalez, Ralf Stumm, Elvira Mass, Elisa Gomez Perdiguero, Anna-Katerina Hadjantonakis, Frederic Geissmann

**Author notes:** Corresponding authors: T. Lazarov and F. Geissmann. Contributed equally (Tomi Lazarov, Pierre-Louis Loyher, Hairu Yang).

## Abstract

Mammalian hematopoietic cells arise from mesodermal progenitors in a close developmental relationship with endothelium, and along three distinct cell lineages known as primitive, pro-definitive, and definitive hematopoiesis. However, the developmental hierarchy between mesodermal progenitors, endothelium, and blood cell lineages is incompletely understood. We report here the identification in mouse gastrula and human (h)iPSC cultures of a population of CXCR4^+^ primitive streak stage mesodermal progenitors that give rise to yolk sac endothelium, blood islands, yolk sac hematopoiesis, and resident macrophages, corresponding to the pro-definitive lineage. Strikingy, this progenitor does not give rise to primitive erythropoiesis or to caudal endothelium and definitive hematopoiesis. Interestingly however, the pro-definitive progenitor population also gives rise to rostral endothelium that persists in adults. Finally, pro-definitive progenitor-derived endothelium is the first and main source of macrophages in embryo and hiPSC cultures, both directly and via previously described multipotent Erythro-Myeloid Progenitors. The identification and isolation of this mesodermal progenitor defines a revised pro-definitive angio-hematopoietic lineage and provides a framework for resident macrophage and endothelial differentiation relevant to disease pathophysiology and their manipulation for therapeutic purposes.

**One sentence summary:** Identification in gastrulating embryo and hiPSC cultures of a mesodermal progenitor for yolk sac and rostral endothelial cells, early macrophages, and Erythro-Myeloid Progenitors characterize pro-definitive hematopoiesis and provide a revised framework for early angio-hematopoietic development.

**Graphical abstract:** Schematic represents the progeny of angio-hematopoietic progenitors at the posterior primitive streak. Developmental timing of CXCR4 expression distinguishes **i)** CXCR4^-^ primitive hematopoiesis, ***ii)*** Early C<u>X</u>CR4^+^ <u>a</u>ngio-hematopoietic <u>p</u>rogenitors (EXAP) (purple) which <u>express</u> CXCR4 between E6.5 and E7.5 and give rise to rostral vessels and resident macrophages through YS hemogenic endothelium, and ***iii)*** late CXCR4^*+*^ intraembryonic angio-hematopoietic progenitors (blue) which acquire expression of CXCR4 around E8 and give rise to definitive hematopoiesis through an intraembryonic hemogenic endothelium. Abbreviations: E (Mouse Embryonic Day); YS (Yolk Sac); Hemog. (Hemogenic); Endo. (Endothelium); Defi. (Definitive).

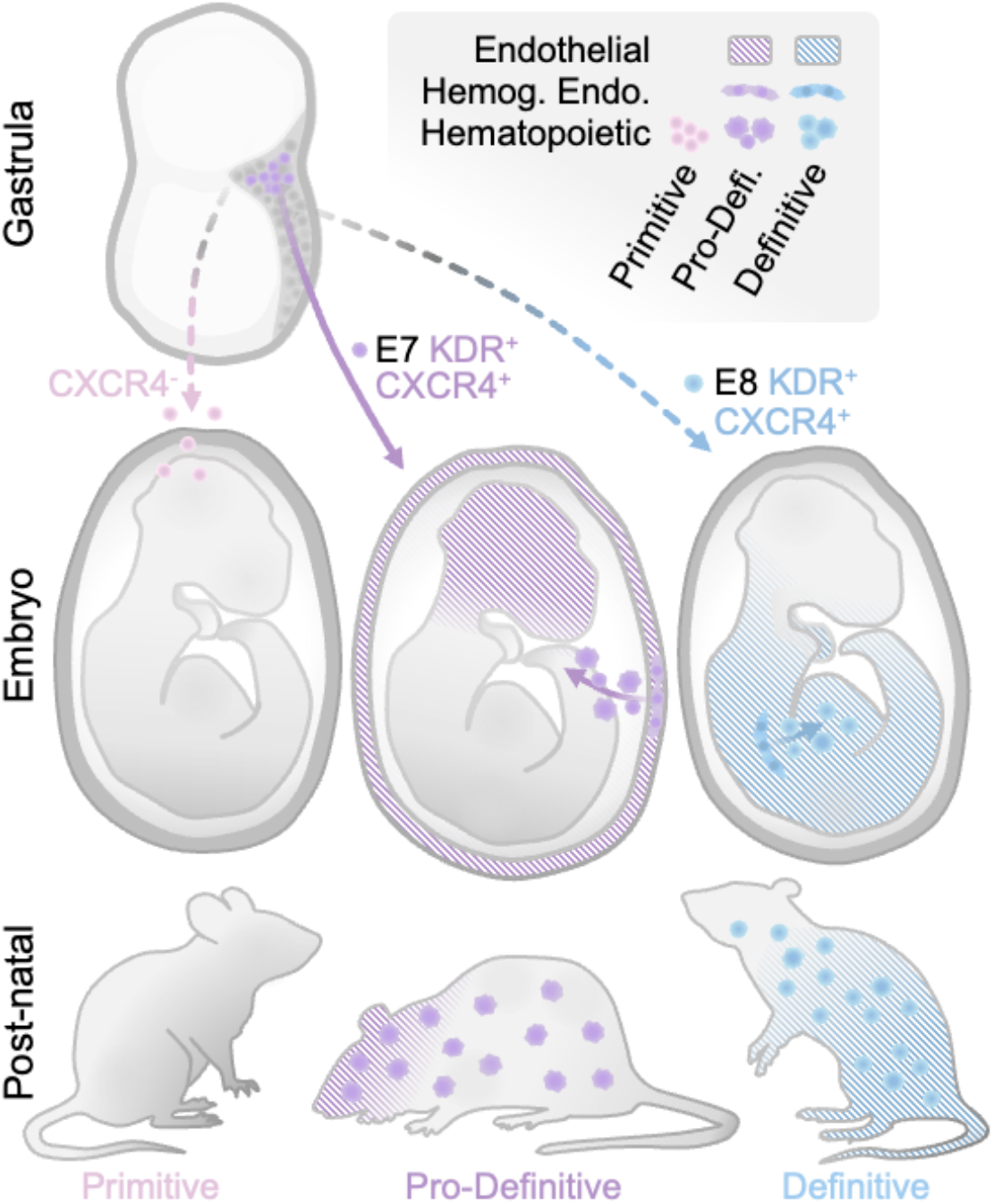

## Introduction

Understanding the development of mammalian hematopoietic and endothelial cells is of crucial relevance for understanding their functions, their roles in diseases, and for advancing regenerative medicine^1,2^. Hematopoietic cells and endothelium arise during embryogenesis in a close developmental relationship^3-7^. A common mesodermal progenitor expressing KDR (FLK1/ VEGFR2) located in the vicinity of the gastrula posterior primitive streak was originally termed angioblast or hemangioblast^3,4,8-13^. However, KDR^+^ angio-hematopoietic progenitors were later found to be heterogeneous^7,14-17^, and seminal studies identified three distinct blood lineages referred as primitive, pro-definitive, and definitive hematopoiesis^1,18-24^, which could not be ascribed to a single common progenitor. At the present time, despite considerable progress in the understanding of hematopoietic development at the cellular and genetic level and at single cell resolution^3,7,14-16,18,20,21,25-53^, the early steps of mammalian angio-hematopoietic remain unclear. For example, at what stage do endothelial(s) and hematopoietic lineages separate, and whether distinct subsets of angio-hematopoietic progenitors are responsible for hematopoietic lineages remain to be demonstrated.

Pro-definitive hematopoiesis gives rise to transient erythrocytes, megakaryocytes, granulocytes, and mast cells that support late embryonic development^18,48,49,54,55^, but also to tissue resident macrophages^49,50,56^ which persists in adults and are essential for tissue homeostasis and repair. Pro-definitive hematopoiesis takes place in the extra-embryonic yolk sac, via a RUNX1-dependent NOTCH1-independent hemogenic endothelium that generates Erythro-Myeloid Progenitors (EMPs)^38,39,54,57-60^ within hemato-endothelial structures known as blood islands^29,40,59-64^. However, the mesodermal progenitor for pro-definitive hematipoiesis and its putative endothelial progeny are otherwise unknown. Furthermore the detection of early macrophage precursor activity^41,43,44,65^ and late macrophage precursor activity^42,46,66^ in embryos suggested that resident macrophages may also originate from the primitive lineage or the definitive lineage. We reasoned that the existence of a genetically tractable committed progenitor for the pro-definitive lineage may therefore help the definition of this lineage and its functions.

In the present studies, we took advantage of the observation that the evolutionary conserved chemokine receptor CXCR4^67-69^, widely expressed among hematopoietic cells including mesodermal KDR+ (FLK1/ VEGFR2) cells^3,70,71^, is regulated in a spatio-temporal manner in different hematopoietic lineages^16,26,72-74^. Time-course lineage tracing, cloning, and transcriptomic and genetic analyses of CXCR4-expressing cells in mouse gastrula and human iPSC cultures, allowed us to identify a CXCR4^+^ primitive streak stage mesodermal progenitor among KDR^+^ cells, which give rise to pro-definitive hematopoiesis via RUNX1-dependent NOTCH1-independent hemogenic endothelial cells in the yolk sac, but not to either primitive erythropoiesis or definitive hematopoiesis. Interestingly, we found that this progenitor also give rise to rostral embryonic endothelial cells. Moreover, these studies also revealed that CXCR4^+^ primitive streak stage mesodermal progenitor -derived yolk sac hemogenic endothelial cells give rise not only to EMPs, but also to the first wave of ‘macrophages-only’ colonies previously attributed to the primitive lineage. We propose that these results therefore define a revised pro-definitive angio-hematopoietic lineage, clarify the origin of adult resident macrophages, and more largely contribute to a revised framework for the developmental hierarchy of the hemato-endothelial system, which should have important implications for future studies aiming to understand the functional diversity of endothelial and hematopoietic cells, the pathophysiology of clonal disorders within these cell lineages, and for devising new strategies to generate cells such as resident macrophages for autologous cell therapies.

## Results

### Primitive/pro-definitive and definitive hematopoiesis arise from spatiotemporally distinct KDR^+^ CXCR4^+^ progenitors

We previously reported that genetic labeling of cells expressing Cxcr4 in *Cxcr4*^*CreERT2*^;*R26*^*LSLtdT*^ mice by administration OH-TAM at embryonic day (E)6.5 resulted in labelling of macrophages but not blood cells, while administration OH-TAM at E9.5 labelled blood cells but not macrophages^74^. We therefore examined single-cell (sc)RNAseq datasets from 6.5 to E8.5 mouse embryos^52^ for expression of Cxcr4 and of Kdr (Vegfr2, Flk1, Cd309), which defines mesodermal angio-hematopoietic progenitors^3,10,11,13,28,75^ (**Fig. 1A, Extended Data Fig. 1 A,B**). Kdr expression is broader that of Cxcr4^+^, but Kdr cells expressing Cxcr4 were predominantly detected in nascent / mixed mesoderm around ∼E7.0 (**Fig. 1A)**, and in PECAM1/VE-CAD^+^ hemato-endothelial progenitor cells after E8.0 (**Fig. 1A, Extended Data Fig. 1 A,B**). Consistently, analysis of staged *Cxcr4*^gfp^ embryos^76^ showed that GFP, which becomes detectable in 4 to 6 hours in mammalian cells^77,78^, was expressed by *Kdr*^+^ cells located in the posterior primitive streak gastrula at the early-bud stage (E∼7.25), previously described as ‘hemangioblasts’^79^ (**Fig. 1B, Extended Data Fig. 1C**), and by *Kdr*^*+*^ cells frequently co-expressing CD31 located within or near the embryo dorsal blood vessels at ∼E8.25 GFP (**Fig. 1B**).

**Figure 1.**
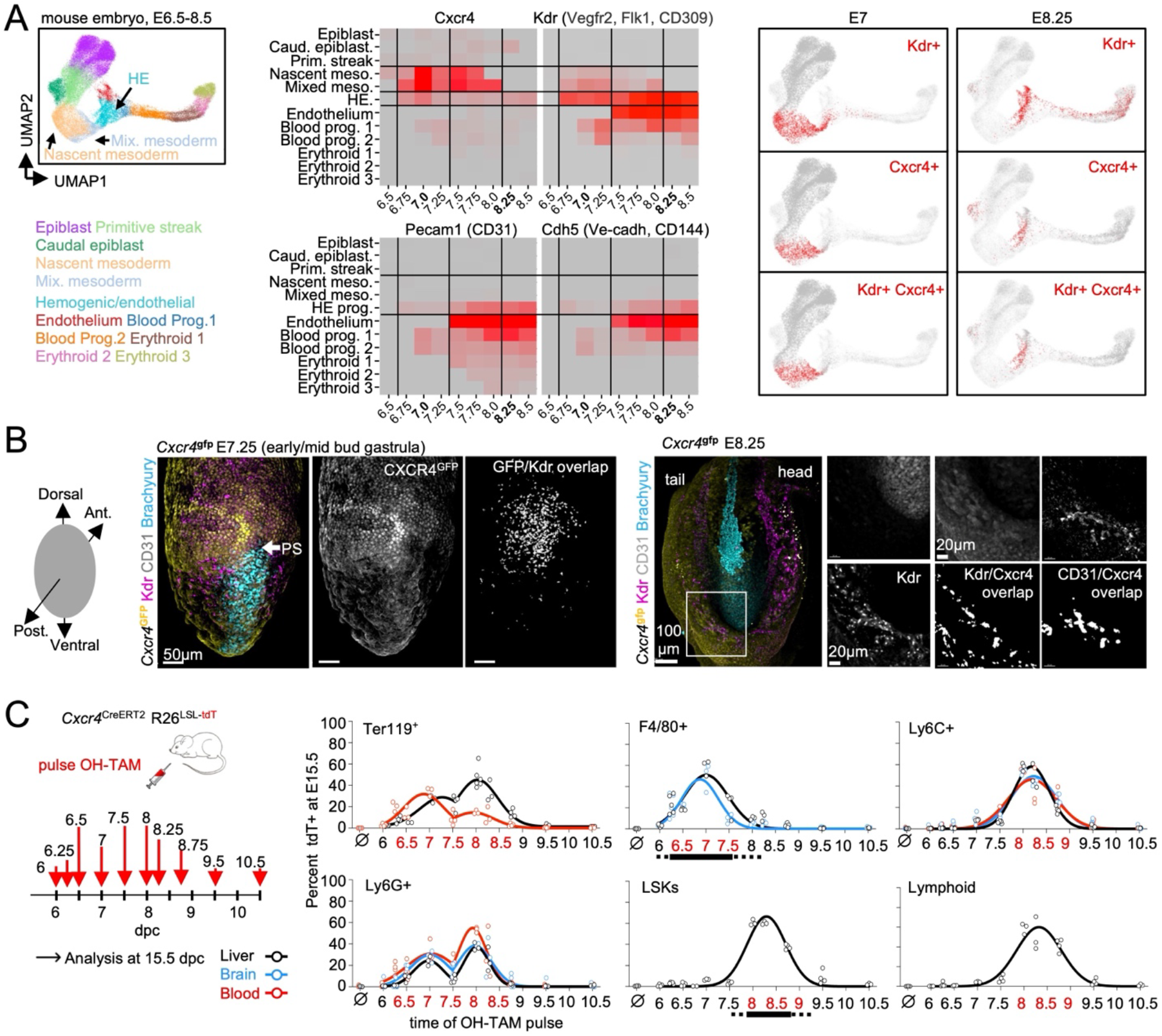
Primitive/pro-definitive and definitive hematopoiesis arise from spatiotemporally distinct KDR+ CXCR4+ progenitors. **A**. Left panel: annotated UMAP scatter plot of single-cell RNAseq analysis of E6.5 to E8.5 dpc mouse embryos^52^, endoderm excluded. Middle panel: Heatmaps show relative expression of select genes in annotated cell types. Right panel: UMAP scatter plots of single-cell RNAseq analysis at E7 and E8.25. Red dots show cells expressing Cxcr4 and/or Kdr at E7 and E8.25. Background grey dots represent all events from E6.5 to E8.5. **B**. Left panel: posterior view of whole-mount immunofluorescence analysis of E7.25 dpc *Cxcr4*^*GFP*^ embryo (early/mid bud stage). Double positive cells were identified by surface-surface statistics (Imaris). PS: primitive streak, white arrow. Representative image from n=3 independent experiments. Right panel: **w**hole mount images of a E8.25 (early somite stage) *Cxcr4*^*GFP*^ embryo at stained with anti-GFP, anti-KDR, anti-Brachyury, anti-CD31 antibodies. Analysis of GFP^+^CD31^+^ and GFP^+^CXCR4^+^ co-expressing cells was performed using Imaris software surface-surface statistics. N=3. **C**. Analysis by flow cytometry of E15.5 *Cxcr4*^*CreERT2*^; *R26*^*LSL-tdT*^ embryo pulsed with OH-TAM at indicated timepoints (red arrows). Plots show percentage of tdT labeled TER119^+^, Ly6G^+^, F480^+^, Lin^-^Sca^+^Kit^+^ (LSK), Ly6G^-^Ly6C^+^, and CD3^+^/CD19^+^/NK1.1^+^ (Lymphoid) cells in liver, brain, and blood. N=3 to 6 embryos/timepoint from 2 to 3 independent experiments. Abbreviations: Embryonic day (E); 4-Hydroxytamoxifen (OH-TAM); Uniform manifold approximation and projection (UMAP); Progenitor (Prog.); Caudal (Caud.); Posterior (Post.); Primitive (Prim.); Mesoderm (Meso.); Hemato-Endothelial (HE).

We thus performed a time course analysis of the hematopoietic progeny at E15.5 of cells that expressed *Cxcr4* during gastrulation (between E6.25 and E9.5^80^) in *Cxcr4*^*CreERT2*^;*R26*^*LSLtdT*^ mice (**Fig. 1C, Extended Data Fig. 1D)**. TER119^+^ red blood cells and Ly6G^+^ granulocytes were labeled by OH-TAM administration between E6.25 and E8.75 (**Fig. 1C)**. In contrast, F4/80^+^ macrophages were mostly labeled during a E6.25 to E7.5 window of time (**Fig. 1C)**, while Lin^-^ Sca1^+^ Kit^+^ (LSK) cells, as well as lymphoid cells and Ly6C^+^ cells were mostly labeled by OH-TAM administration between E8 and E9 (**Fig. 1C**). Because CreER-mediated reporter expression is detectable 6 to 12 hours after intraperitoneal injection of tamoxifen^81,82^, these data indicated that progenitors for macrophages and for a first wave of red blood cells and granulocytes (primitive and or pro-definitive hematopoiesis) expressed *Cxcr4* between ∼E6.5 and E7.75, while progenitors for LSK, lymphoid cells, Ly6C cells and a second wave of red blood cells and granulocytes (definitive hematopoiesis) expressed *Cxcr4* between E8 and E9.

These studies suggested that a population of Kdr^+^ Cxcr4^+^ early/mixed mesoderm located near the gastrula posterior primitive streak, likely correspond to a subset of previously described hemangioblasts^3,4,8-12^, are committed progenitors for primitive and/or pro-definitive hematopoiesis, and are distinct from progenitors for LSK and lymphoid cells which acquire Cxcr4 expression after E8, near or within intraembryonic dorsal blood vessels.

### Early Cxcr4+ progenitors give rise to pro-definitive hematopoiesis and rostral blood vessels

To better characterize the angio-hematopoietic progeny of the early (∼E7) CXCR4^+^ progenitors, we conducted a time-course analysis of the fate of progenitors pulsed with OH-TAM at E6.5. *Kdr*^+^ cells were labeled at E7.5 and E8.5 (**Fig. 2A, Extended Data Fig. 2A**). CD31^+^ endothelial cells and phenotypic erythro-myeloid progenitors (EMPs, CD41^+^ CD93^+^ Sca1^-^ Kit^+^^18,83-85^) were labeled from E8.5 as they become detectable, first in the yolk sac (YS) (**Fig. 2A-C**), where they formed blood islands^40^, with CD31^+^ endothelial cells and adjacent CD41^+^ hematopoietic cells^39^ (**Fig. 2B**). At later time points, 9.5, 10.5, and 12.5, intra-embryonic endothelial cells were labeled, with decreasing efficiency along the rostro-caudal axis (**Fig. 2C,D, Extended Data Fig. 2B**). For example, cephalic blood vessels were labeled (**Fig. 2C to E**) while the dorsal aorta (**Fig. 2E**) and caudal vessels in general (**Fig. 2C,D**) were not. F4/80^+^ macrophages were also labeled in the YS, rostral, and caudal embryo as they become detectable from E9.5 onwards^65,86^ (**Fig. 2C**). Analysis of 4 weeks-old mice confirmed labeling of cephalic vessels and tissue macrophages^48,49,74^ (**Fig. 2F**).

**Figure 2.**
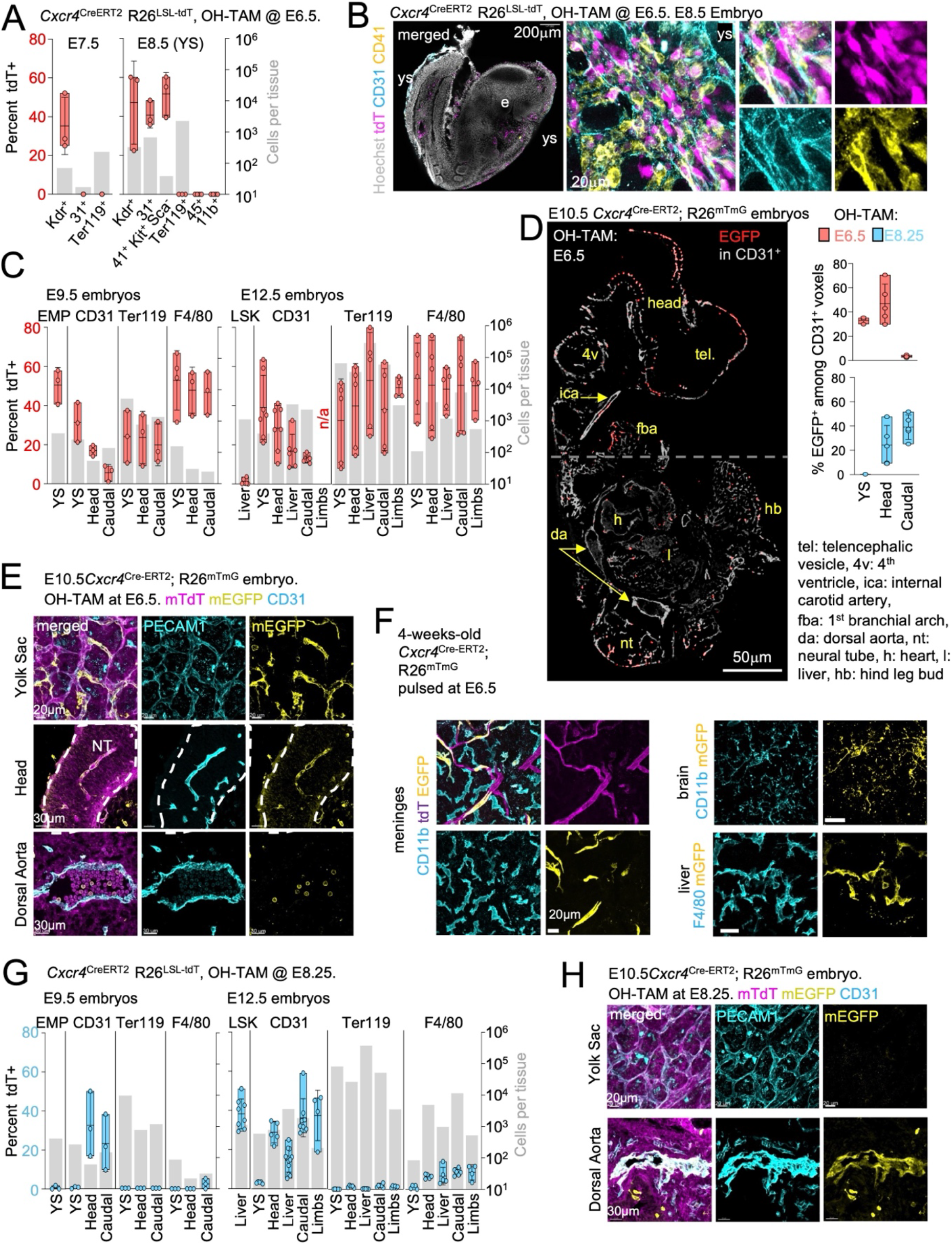
Early Cxcr4+ progenitors give rise to pro-definitive hematopoiesis and rostral blood vessels. **A**. *Cxcr4*^*CreERT2*^; *R26*^*LSL-tdT*^ embryos treated with OH-TAM at E6.5 were analyzed by flow cytometry at E7.5 and E8.5. Red axis: % tdT labeled cells and grey axis: total cell number. N=3 to 5 embryos/timepoint. **B**. Whole mount *Cxcr4*^*CreERT2*^; *R26*^*LSL-tdT*^ embryos treated with OH-TAM at E6.5 were analyzed at E8.5 by immunofluorescence. Representative images from n=3 independent experiments **C**. *Cxcr4*^*CreERT2*^; *R26*^*LSL-tdT*^ embryos treated with OH-TAM at E6.5 were analyzed at E9.5 and E 12.5 by flow cytometry. Red axis: % tdT labeled cells and grey axis: total cell number in indicated organs and cell populations. EMP: CD41^+^ CD93^+^ Sca1^-^ Kit^+^, LSK: Lin^-^ Sca1^+^ Kit^+^. n/a: not available. N=3 to 7 embryos/timepoint from 2 to 4 independent experiments. **D,E**. Wholemount (YS) or cryosections (embryo proper) from E10.5 (Kaufman TS 16) *Cxcr4*^*CreERT2*^; *R26*^*mTmG*^ embryos treated with OH-TAM at 6.5 dpc were analyzed by immunofluorescence. In D left panel displays mEGFP+ voxels (red) among CD31^+^ voxels in sagittal section. Gray dashed line indicates the border between the head and caudal region. Right panel plots quantify % of mEGFP+ voxels among CD31+ voxels in the YS, head, or caudal regions of 10.5 dpc embryos from mice treated with OH-TAM at E6.5 and E8.25 for comparison. n= 3-6 embryos per timepoint, from 3 independent experiments. **F**. 4-week-old *Cxcr4*^*CreERT2*^; *R26*^*mTmG*^ mice treated with OH-TAM at E6.5: whole-mount meninges, brain parenchyma and liver were analyzed by immunofluorescence. **G**. E9.5 and E12.5 *Cxcr4*^*CreERT2*^; *R26*^*LSL-tdT*^ embryos treated with OH-TAM at E8.25 were analyzed by flow cytometry as in (C). Blue axis: % tdT labeled cells and grey axis: total cell number. N=3 to 8 embryos/timepoint from 2 to 4 independent experiments **H**. Cryosections from E10.5 *Cxcr4*^*CreERT2*^; *R26*^*mTmG*^ treated with OH-TAM at E8.25 were analyzed by immunofluorescence. n>3 mice per timepoint, from at least 3 independent experiments.

Interestingly, although Ter119^+^ red blood cells were detectable from E7.5 and abundant in the E8.5 yolk sac^43,61,62^ they were not labeled by administration of OH-TAM at E6.5 (**Fig. 2A**). The labeling of TER119^+^ erythroid cells became detectable at E9.5 and increased at 12.5 (**Fig. 2C**). An earlier administration of OH-TAM at E 5.75 also failed to label E8.5 Ter119^+^ red blood cells (**Extended Data Fig. 2C)**. In addition, analysis of genes differentially expressed in Kdr^+^Cxcr4^+^ cells in comparison to Kdr^+^Cxcr4^-^ cells at E7 indicated lower expression levels of the homeobox *Mixl1*, which supports primitive erythroid differentiation from the primitive streak^87,88^ and *Id1* which restricts myeloid commitment, and higher levels of the homeobox *Lhx1*, the endothelial/ hematopoietic transcription factors *Lmo1, Gata4 and Gata6*^89-93^, the endothelial factor *Mesp1*^94^, and *Eomes*, which is important for the hemogenic competence of yolk-sac mesodermal progenitors^34^ (**Extended Data Fig. 1A)**. These data suggest that the ∼E7 CXCR4^+^ progenitors are the source of pro-definitive hematopoiesis rather than of primitive hematopoiesis.

In contrast, the later administration of OH-TAM at E8.25 resulted in the absence of labeling in the yolk sac at E9.5, E10.5, and E12.5 (**Fig. 2D,G**), while endothelial cells were labeled predominantly in the cephalic and caudal embryo, including dorsal vessels (**Fig. 2H**), and LSK were labeled in the fetal liver (**Fig. 2G**). Few macrophages were labeled at this stage (**Fig. 2G**). These data, within the limitations of the murine model in terms of lineage specificity^95^ and for the clonal analysis of precursor-products relationships, identify a dedicated subset of mesodermal Kdr^+^ ‘hemangioblasts’ characterized by Cxcr4 expression at the primitive streak at E7, or <u>e</u>arly <u>X</u>4 <u>angio-hematopoietic p</u>rogenitor (EXAP), which give rise to yolk sac and rostral endothelial cells, and yolk sac pro-definitive hematopoiesis (blood islands, Sca1^-^ hematopoietic precursors, yolk-sac derived macrophages, granulocytes and erythroid cells arising after E8.5). In contrast, EXAP does not contribute to the first wave of erythroid cells -corresponding to primitive erythropoiesis^43,70^-, and to caudal blood vessels including the dorsal aorta, and the LSK/definitive hematopoietic lineage, which arise from a progenitor that do not express Cxcr4 before E8.

### CXCR4^+^ KDR^+^ angio-hematopoietic progenitors in developing human embryos and hiPSC cultures

Considering the above limitations and the known differences in yolk sac and early hematopoietic development in human and mice^96^, we next sought to address whether the above findings in our studies of mouse development are conserved and can be extended to human. We identified cells co-expressing *KDR* and *CXCR4* among populations annotated as nascent and mixed mesoderm, yolk sac mesoderm, and extra-embryonic hemogenic endothelial progenitors in a single-cell (sc)RNAseq dataset of a Carnegie stage 7 human embryo^53^ which could therefore represent a candidate human EXAP (**Extended Data Fig. 3A**). We therefore investigated the existence and fate of putative CXCR4^+^ progenitors in a human induced-pluripotent stem cell (hiPSC) model, tractable at the genetic and cellular levels, that recapitulates aspects of yolk-sac hematopoiesis^97-99^ (**Extended Data Fig. 3B to F**). Time-course analysis of the phenotype, morphology, and numbers of cells arising in cultures from 3 independent hiPSC lines identified putative KDR^+^ Lin^-^ and KDR^+^ CXCR4^+^ precursors, early CD45^-^ CD235a^bright^ erythroid cells, CD144^+^ endothelial cells, and CD45^+^ hematopoietic cells (**Fig. 3A, Extended Data Fig. 3B-E**). On average, 1/3^rd^ of cells expressed KDR (Pop.1) at the onset of hiPSC culture as reported for ES cells^14^ **(Fig. 3B)**. The first 12 hours of culture were marked by early expansion of small cells co-expressing KDR and CXCR4 (Pop. 2), of larger CXCR4^-^ erythroid cells expressing high level of Glycophorin A (CD235a^bright^, Pop. 3), and of cells expressing the endothelial marker VE-Cadherin (CD144, Pop.4) (**Fig. 3B**). CD144^+^ cells were heterogeneous in size with minor subsets expressing the hematopoietic antigens CD235, CD43, or CD41 after ∼1 day of culture (**Fig. 3A)**.

**Figure 3.**
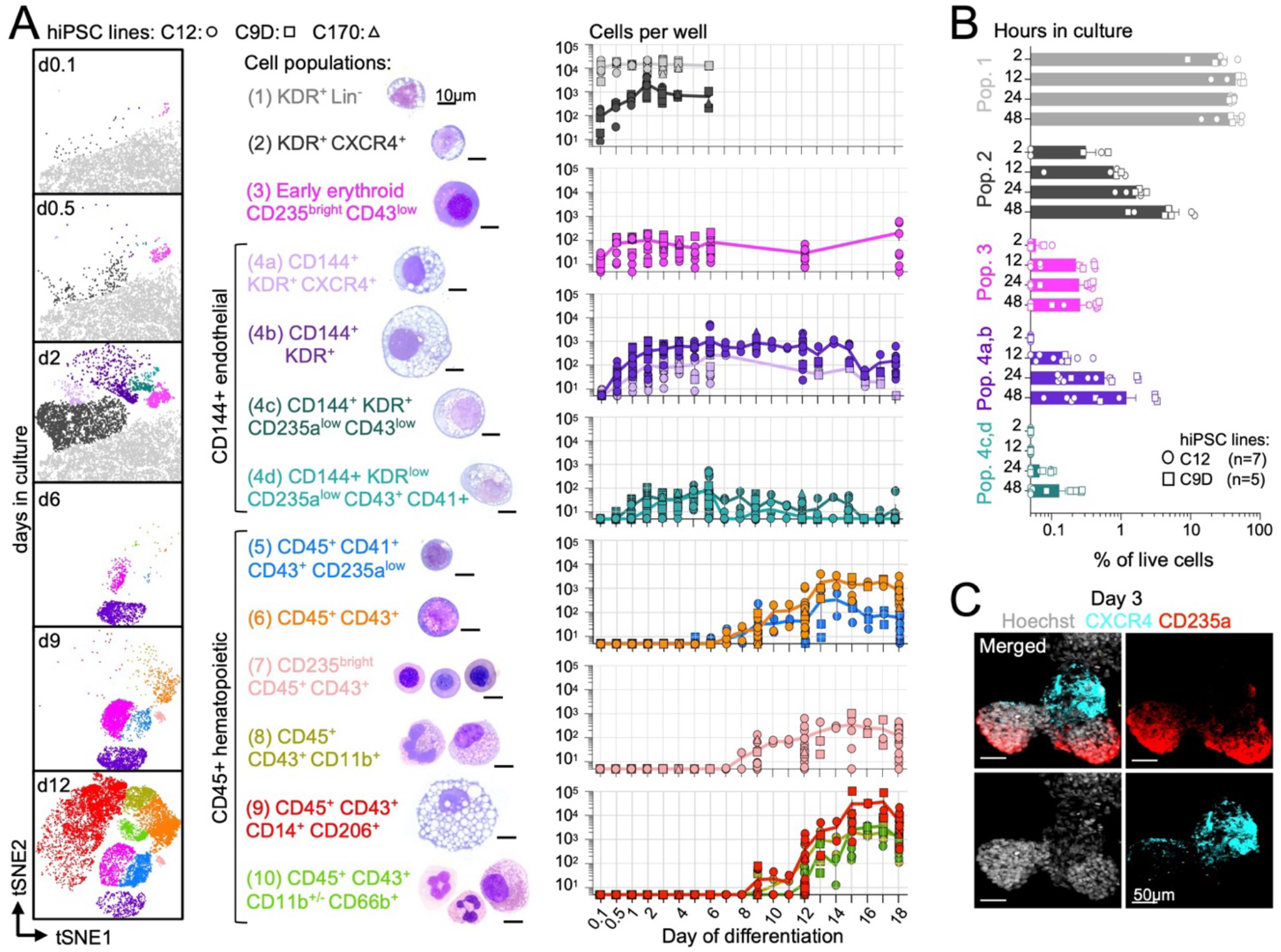
Human CXCR4^+^ cells in hiPSC cultures. **A**. hiPSC differentiation cultures. Left: tSNE representation of flow cytometry analysis, color-coded by cell phenotype. CXCR4^+^ and KDR^+^ cells are not shown on day 6 to day 12 plots. Middle: phenotype and representative May-Grunwald Giemsa staining for cell populations. Right: Cell number per well over time. n: 3 independent cell lines and 2 independent experiments per line. **B**. Percent of indicated cells among total live cells during the first 48hrs of differentiation. N=5 and 7 experiments with 2 cell lines. **C**. Whole-mount immunofluorescence imaging of day 3 EBs. Representative from 3 independent experiments.

CD45+ hematopoietic cells become detectable around day 6 (**Fig. 3A)**. We did not obtain lymphoid cells in these culture conditions **(Extended Data Fig. 3F)**. Of note cells expressing CD235^bright^ or CXCR4 were spatially segregated in early EBs (**Fig. 3C**). These data indicate the presence of KDR^+^ CXCR4^-^ and KDR^+^ CXCR4^+^ progenitors at the onset of human angio-hematopoietic differentiation. They also suggest independent development of early erythroid cells and of CXCR4^+^ cells.

### Expression of CXCR4 biases the differentiation potential of hiPSC towards pro-definitive hematopoiesis

Clonal analysis of the hematopoietic and endothelial differentiation of Facs-purified KDR^+^ (Pop.1) cells on OP9 stroma (**Fig. 4A, Extended Data Fig. 4**) indicated that 1/3^rd^ of positive wells give rise to pure CD235a^bright^ erythroid cells, ∼40% to pure endothelial CD144^+^ cells, and ∼25% to endothelial cells mixed with either CD14^+^ macrophages, CD235a^bright^ erythroid cells, or both (**Fig. 4A**). In contrast, KDR^+^ CXCR4^+^ (Pop.2) cells did not give rise to erythroid-only colonies (**Fig. 4A**), but to pure endothelial cells in 60% of positive wells, endothelial cells mixed with macrophages in ∼25% of wells, endothelial cells mixed with erythroid cells in ∼10% of wells, or endothelial cells mixed with both erythroid and macrophage in ∼5 % of wells (**Fig. 4A**). Bulk cultures of purified populations confirmed that Pop.3 differentiate from Pop.1 progenitors but not from Pop.2 after culture on OP9 stroma (**Fig. 4B, Extended Data Fig. 4B,C**). Of note, FACS-purified KDR^-^ CXCR4^+^ cells did not give rise to endothelial or hematopoietic progeny (**Extended Data Fig. 4D**). Analysis of RUNX1-deficient and isogenic hiPSC lines (**Extended Data Fig. 5, S6)** showed that differentiation of Pop.3 was RUNX1-independent (**Fig. 4C, Extended Data Fig. 6**), indicating it does not require endothelial to hematopoietic transition (EHT) as expected for primitive erythrocytes^25,57-60^. Moreover, Pop.3 gave rise to erythroid only colonies on OP9 stroma and Methocult (**Fig. 4D, Extended Data Fig. 4E)**, independently of RUNX1 (**Fig. 4D**), and expressed hemoglobin epsilon (HBE) (**Fig. 4E**) These data showed that KDR^+^ cells (Pop. 1) have the capacity to differentiate into primitive erythroid progenitors (Pop.3), and into KDR^+^ CXCR4^+^ cells (Pop.2), the latter lacking primitive erythroid potential.

**Figure 4.**
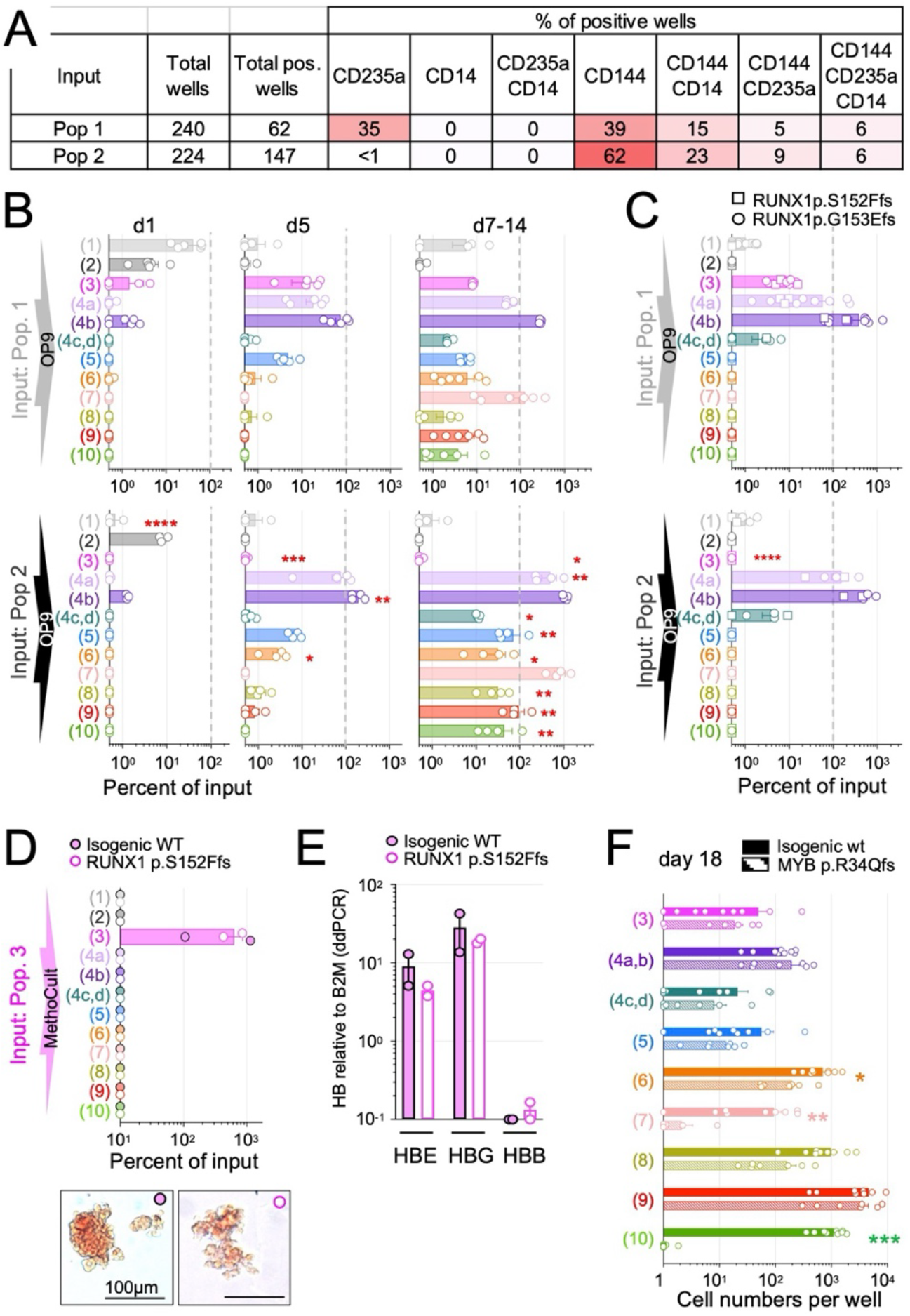
Angio-hematopoietic potential of human CXCR4^+^ and CXCR4^-^ progenitors. **A**. Cloning analysis: FACS-purified cells from Pop.1 and 2 plated at estimated density of 0 or 1 cell per well, cultured for 14 days on OP9 stroma, and wells are stained for immunofluorescence analysis. Results from 2 independent experiments. **B**. 10^3^ FACS-purified Pop.1 and 2 cells cultured for 1, 5, 7, or 14 days on OP9 stroma and their progeny analyzed by flow cytometry. Cell numbers plotted as percentage of input. p-values between Pop.1 and 2 progeny calculated by Mann-Whitney test. n= 3 to 6 independent experiments/timepoint. **C**. 10^3^ FACS-purified isogenic WT or RUNX1-null (p.S152Ffs and p.G153Efs) Pop.1 and 2 cells analyzed at d7 as in (B). **D**. 10^3^ FACS-purified cells from isogenic WT or p.S152Ffs Pop.3 cultured for 14 days in MethoCult are analyzed by flow cytometry and brightfield microscopy. n=2 independent experiments by genetic background. **E**. HBE, HBG, and HBE mRNA expression by ddPCR by isogenic WT or p.S152Ffs FACS-purified Pop.3 cells. n=2 experiments per genotype. **F**. EBs from isogenic WT and MYB-null (p.R34Qfs) hiPSCs analyzed by flow cytometry at day 18, n=7 independent experiments. Statistics: Mann-Whitney test, p≤0.05 (*); p≤0.01 (**); p≤0.001 (***); p<0.0001 (****)

In contrast, Pop.2 cells generated ∼10 times more endothelial and late hematopoietic cells than Pop.1 (**Fig. 4B**), and RUNX1 was required for their differentiation into CD45^+^ hematopoietic cells (pop 5 to 10) (**Fig. 4C, Extended Data Fig. 6**). In addition, analysis of MYB-deficient and isogenic hiPSCs (**Extended Data Fig. 5)** showed that differentiation of endothelial cells and macrophages was not affected by MYB deficiency, while hematopoietic precursors, erythroid cells, and granulocytes were either decreased or missing (**Fig. 4F, Extended Data Fig. 6B**). The transcription factor MYB is also dispensable for differentiation of yolk-sac derived macrophages in mice^49^, while necessary for differentiation/ survival of yolk-sac derived multipotent hematopoietic progenitors, late erythroid cells, and granulocytes (**Extended Data Fig. 7**) as well as for LSK-derived definitive hematopoiesis^49,66,100,101^. These data altogether indicate that KDR^+^ CXCR4^+^ cells (Pop.2) behave like a human EXAP, which does not give rise to RUNX-1 independent primitive erythroid cells, but generate endothelial cells, either alone in 2/3^rd^ of wells or associated with RUNX1-dependent hematopoietic cells in ∼1/3^rd^ of wells, which resemble yolk sac-derived RUNX1 dependent pro-definitive hematopoiesis in mice, including MYB-dependent erythroid cells and granulocytes and MYB-independent macrophages.

### hEXAP generates MYB-independent macrophages via NOTCH1-independent blood islands

The above data are compatible with the hypothesis that RUNX1-dependent late hematopoietic cells arise from Pop.2 via a hemogenic endothelial intermediate^39^. Clonal analysis of the progeny of CD144^+^ cells cultured on OP9 stroma indicated that ∼70% of wells gave rise to endothelial cells alone, 15% to endothelial cells mixed with macrophages, ∼10% to endothelial cells mixed with erythroid cells and ∼5% to endothelial cells mixed with both erythroid cells and macrophages **(Fig. 5A**). These frequencies were comparable with the clonal analysis of Pop.2 cells (see Fig. 4A). Analysis of FACS-purified bulk Pop. 4a, 4b, 4c, and 4d further showed that endothelial subset gave rise to each other and to CD45^+^ hematopoietic cells (**Fig. 5B)**, in a RUNX1 dependent manner **(Extended Data Fig. 8 A to C)**. Interestingly, the ratio of CD45^+^ hematopoietic to endothelial progeny increased from 0.3 (pop.4a), to 0.6 (pop.4b), 3 (pop. 4c) and to 20 (pop. 4d) (**Fig. 5C**), the latter demonstrating a strong hematopoietic bias associated with the co-expression of hematopoietic antigens (CD43, CD41, and CD235a) by endothelial cells. Imaging studies further illustrated the progressive changes of the differentiation potential of CD144^+^ subsets (**Fig. 5D**), from a large network of CD144^+^ cells occasionally containing hematopoietic CD41^+^ clusters^39^ reminiscent of “endothelial blood island”-type structures (Pop.4a), to the predominance of CD144^+^/CD41^+^ blood island”-type clusters (Pop.4c and d). Finally, Pop.4a, b, c, and d developed and differentiated in the absence of NOTCH1 (**Fig. 5E, Extended Data Fig. 5, S6B**) as observed for yolk sac hemogenic endothelium in mice^27,33,102,103^, while the hemogenic endothelium which give rise to LSKs is NOTCH1-dependent^27,102^. To understand whether large KDR^+^ CD144^+^ endothelial cells (Pop. 4b) from late EB can still be induced to produce hemogenic endothelium and hematopoietic cells, they were Facs-purified at day 16 of culture and re-cultured on OP9 stroma. In this setting, Pop. 4b cells also generated Pop.4a, c, d and hematopoietic cells (**Extended Data Fig. 8D**). These data therefore suggest that hEXAP (Pop.2) generates endothelial cells that mature into NOTCH1-independent ‘yolk sac like’ hemogenic endothelium, form blood-island like structures, and produce CD45+ hematopoietic cells, including MYB-independent macrophages.

**Figure 5.**
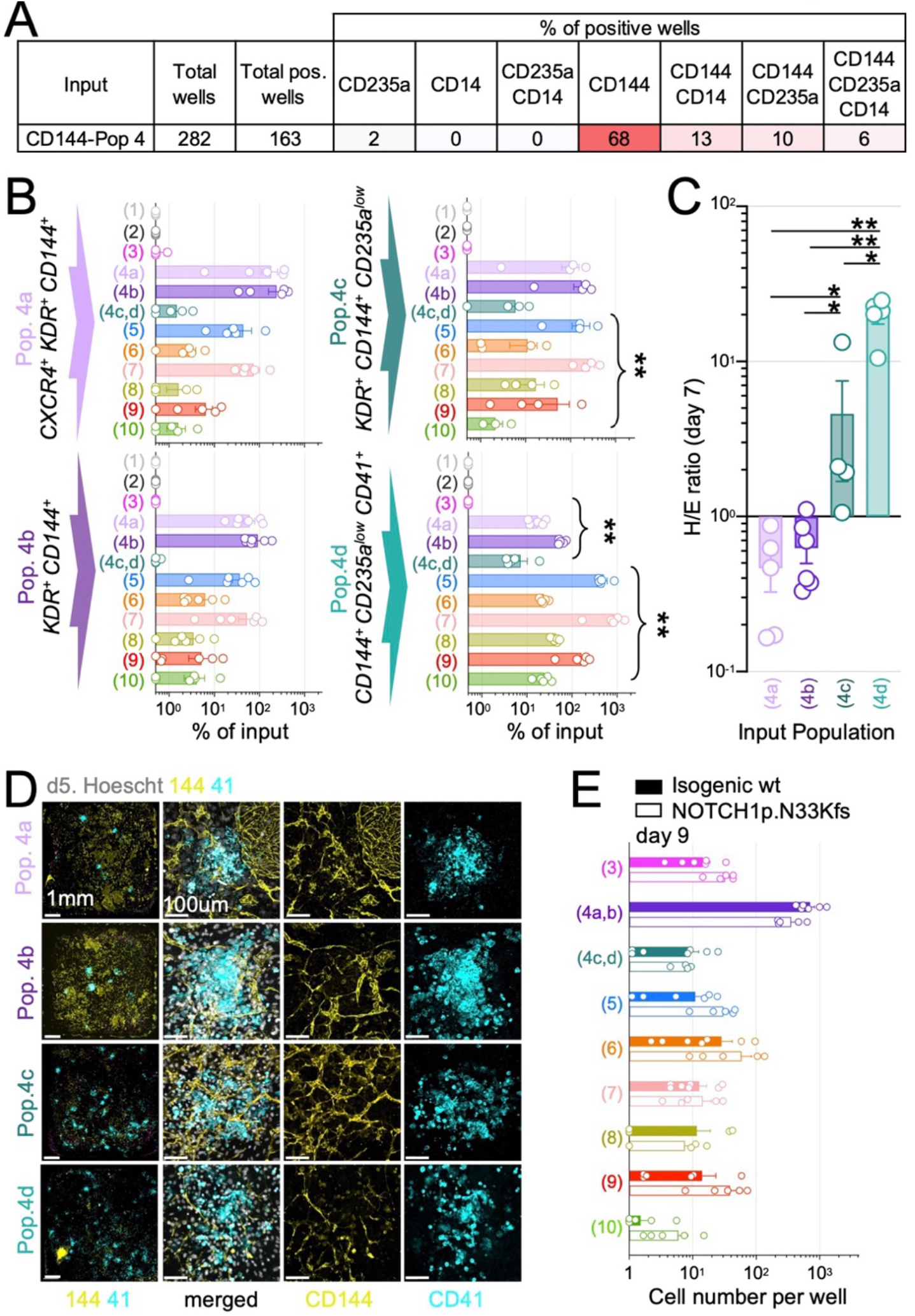
hEXAP generates MYB-independent macrophages via NOTCH1-independent blood islands. **A, B**. Progeny of FACS-purified cells from Pop.4a, b, c, and d analyzed as in (4A). Results from 2 independent experiments. **B**. Progeny of FACS-purified cells from Pop.4a, b, c, and d analyzed as in (4B), p-values between Pop.1 and 2 progeny calculated by Mann-Whitney test. n= 3 to 6 independent experiments/timepoint. **C**. Ratio of hematopoietic (sum of Pops 3 and 5 to 10) to endothelial (sum of Pops 4a-d) output from experiments as shown in B. Statistical comparisons by Mann-Whitney test. Results from 5 independent experiments. **D**. 5×10^2^ live FACS-purified 4a, 4b, 4c, or 4d cells cultured for 5 days on OP9 stroma were analyzed by immunofluorescence, representative images of 3 independent experiments. **E**. Differentiation of Isogenic WT and NOTCH1-null (p.N33Kfs) iPSC lines analyzed by flow cytometry at day 9, cells numbers per well. Results from 5 independent experiments. P-values: ns by Mann-Whitney test. p≤0.05 (*); p≤0.01 (**); p≤0.001 (***); p<0.0001 (****).

### Early macrophage potential of EXAP-derived hemogenic endothelium

As noted above, clonal analysis indicated that EXAP and CD144+ cells give rise to both erythroid and myeloid cells at the population level, but that macrophages and erythroid progenies unfrequently arise from the same precursors at the clonal level (see Fig. 4A and Fig. 5A). Analysis of the differentiation potential CD45^+^ cells in MethoCult and OP9 stroma (**Fig. 6A,B, Extended Data Fig. 8E**) confirmed that although phenotypic EMPs (Pop.5, CD45^+^ CD43^+^ CD41^+^ CD235a^low18,63,104^) give rise to both erythroid and myeloid cells at the population level (**Fig. 6A**), separate erythroid and myeloid colonies are frequently observed in MethoCult (**Fig. 6B**) as routinely observed in mice^18,48,63^. Pop.6 (CD45^+^ CD43^+^ CD41^-^ CD235a^-^) and Pop. 7 (CD45^+^ CD43^+^ CD41^-^ CD235^bright^) arise around the same time as Pop.5 in hiPSC differentiation cultures (see Fig. 3A) and proliferate to generate either myeloid or erythroid colonies respectively (**Fig. 6C**). Population 8, 9 and 10 were weakly proliferative (**Extended Data Fig. 8E)**. These data suggest that KDR^+^ CXCR4^+^ progenitors may generate endothelial cells with a myeloid or erythroid hemogenic bias, as well as a bipotent erythro-myeloid potential. These findings are consistent with previous work showing that mouse blood islands are polyclonal^7,29^, and also with the detection of macrophage precursor in the mouse E∼7 embryo^41,43,44^, at the time of primitive erythropoiesis and before the emergence of erythro-myeloid progenitors, which was interpreted at the time as the existence of primitive macrophages.

**Figure 6.**
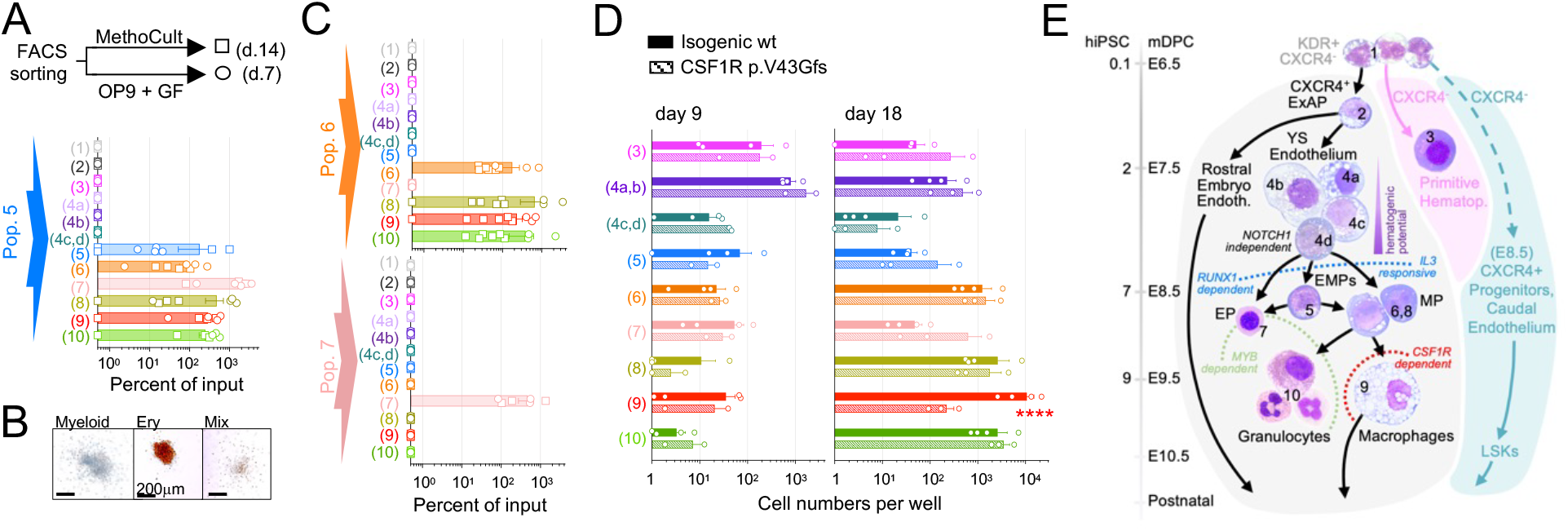
EXAP-derived CD45+ hematopoietic cells. **A,B**. FACS-purified Pop.5 (5×10^2^), cultured for 7 days on OP9 stroma (circles) or 14 days in MethoCult (squares) and analyzed by flow cytometry (A) and brightfield microscopy (B). Micrographs are representative MethoCult colonies. n = 6 independent experiments. **C**. As in (A) for Facs purified Pop.6 (5×10^2^) and Pop.7 (5×10^2^) cells. **D**. EBs from isogenic WT and CSF1R-null (p.V43Gfs*20) hiPSCs analyzed by flow cytometry at day 9, n=4 and 2 independent experiments, and d18 n=4 and 3 independent experiments. Statistics: Mann-Whitney test, p≤0.05 (*); p≤0.01 (**); p≤0.001 (***); p<0.0001 (****). **E**. Schematic represents an interpretation of the results depicted in Figs 1 to 5.

To further investigate the existence of early CSF1R-dependent macrophage progenitors, we analyzed differentiation of isogenic and CSF1R-deficient hiPSCs (**Extended Data Fig. 5, S6B**). Results showed no difference in endothelial and hematopoietic/macrophage differentiation at early time points (**Fig. 6D**). However, macrophage number was selectively reduced by CSF1R-deficiency at late time points (**Fig. 6D**), compatible with the important role of CSF1R for macrophage survival. Altogether, these results show that mouse and human KDR^+^ CXCR4^+^ progenitors give rise to RUNX1-dependent / NOTCH1-independent hemogenic endothelial cells, organized in blood islands, which generate hematopoietic precursors endowed with a variety of myeloid, erythroid, and mixed erythro-myeloid fates, including MYB-independent macrophages in hiPSC cultures and in mice, while primitive hematopoiesis and LSK originate from distinct progenitors and via different mechanisms (**Fig. 6E**).

## Discussion

The developmental origin of hematopoietic lineages and their relationship with endothelial cells has been obscured by the heterogeneity of putative progenitors, the existence of several hematopoietic and -likely-endothelial waves, and interspecies differences^14-16,25,29^. Our present studies identify a lineage-restricted progenitor for a revised pro-definitive angio-hematopoietic lineage, distinct from primitive erythropoiesis, and from caudal endothelial cells and the LSK lineage.

This population of progenitors, which could be named EXAP for <u>e</u>arly C<u>X</u>CR4 <u>a</u>ngio-hematopoietic <u>p</u>rogenitors, is present at the anatomical site and time where hemangioblasts were initially identified, the primitive streak^9^ at ∼E7.0 in the mouse embryo, human Carnegie Stage 7, and the initial stages of hiPSC differentiation^10^. We show that it gives rise to YS endothelial cells as well as to rostral/cephalic endothelial cells, and to yolk sac hemogenic endothelial cells organized in blood islands which in turn generate RUNX1-dependent pro-definitive hematopoiesis including MYB-independent resident macrophages. In addition, our studies strongly suggest that the detection of macrophage progenitor activity at the time of primitive erythroid cell emergence in mice^41,43,44^, which has been considered evidence for existence of mammalian primitive macrophages, may correspond in fact to EXAP-derived endothelial cell giving rise to macrophage colonies without an EMP intermediate, while the concept of hemogenic endothelium-derived yolk sac EMP^18^ applies at the population level.

Of note the present study should help resolve apparent controversies/uncertainties on the development of macrophages from different hematopoietic or angio-hematopoietic waves^45,47,50^. Our results suggest a conserved developmental origin of the yolk-sac macrophage lineage in mammals that differs from Zebrafish^42,105-107^. In regard to this interspecies difference, it is conceivable that differences in the first steps of hematopoiesis between fish, where primitive granulocytes and macrophages are abundant^106^, and mammals where primitive red blood cells are predominant^18^, may be linked in part to environmental differences, *i*.*e*. the easier access of oxygen and pathogens to the fish embryo.

Our results are also consistent with a developmental topology of the angio-hematopoietic system characterized by orderly anatomical allocation of mesodermal progenitors to distinct lineages. *In vivo* analysis suggests that intraembryonic caudal vessels and definitive hematopoiesis arise from distinct progenitors independent from EXAP and expressing CXCR4 later, around E8 *in vivo* in mouse, consistent with well documented evidence that hemogenic endothelium in the aorta-gonad-mesonephros (AGM) is a source of LSK cells in embryos and adult ^19,22,108^. This late CXCR4^+^ progenitor may represent the *in vivo* counterpart to a recently described angio-hematopoietic progenitor in human iPSC cultures ^16^. In addition, the conclusion that primitive erythropoiesis develops from a distinct progenitor and without a hemogenic endothelial cell intermediate in mammals is also consistent with previous studies^7,15,21^. Altogether, our results are consistent with the existence of -at least-three distinct populations of mesodermal progenitors that give rise successively to primitive hematopoiesis, to pro-definitive hematopoiesis, and finally to caudal vessels and definitive hematopoiesis (see graphical abstract and Fig. 6E).

The identification of a pro-definitive angio-hematopoietic progenitor has wide ranging implications for studies investigating the biology of endothelial and hematopoietic cells. For example, the insights, tools, and methods presented in this work, can serve as a framework to decipher the molecular mechanisms of specification, and investigate the function of developmentally distinct angio-hematopoietic endothelial and hematopoietic cells. Given that our results show commitment towards the pro-definitive lineage occurs at the level of early mesodermal progenitors, our study is also of interest to investigators developing iPSC differentiation methods to derive autologous cells for therapeutic purposes. Our findings are also relevant for the pathophysiology of certain somatic mosaicism associated disorders. We previously demonstrated that somatic mutations arising during development in yolk-sac progenitors of tissue-resident macrophages causes neurodegeneration in mice^109^. Recent works show this mechanism of pathogenesis may contribute to the development and progression of Alzheimer’s disease in human^110,111^. Our present results may inform future studies in this domain, for example, whether somatic mutations in cells from the EXAP lineage during development may contribute to clonal endothelial/macrophage proliferations observed in cerebral cavernomas and myeloproliferative neoplasms^112,113^.

## Acknowledgements

The authors are indebted to our colleagues Alexander Rudensky and Alexander Gitlin for critical reading of the manuscript, and members of the Geissmann lab for their support and helpful suggestions.

## Funding

MSKCC and the MSKCC Stem Cell Core are supported by NIH P30CA008748. Work in FG lab was supported by NIH grants R01NS115715-01, R01 HL138090-01, R01 AI130345-01. Work in AKH was supported by NIH grants R01DK127821, R01HD086478, and P30CA008748. EGP was supported by the Institut Pasteur, the CNRS, Revive (Investissement d’Avenir; ANR-10-LABX-0073) and the European Research Council ERC investigator award (2016-StG-715320).

## Author contributions

FG and TL designed the overall study, analyzed data, and wrote the draft manuscript. TL, HY and TZ generated and characterized mutant iPSC lines. TL designed, performed and analyzed flow cytometry experiments with hiPSC lines, with help from HY. PLL and TL designed, performed, and analyzed imaging experiments with hiPSC lines. PLL designed, performed, and analyzed imaging experiments with mouse embryos with assistance from TL, SN, YYK, AKH and EGP. HY, PLL, ZD and TL designed, performed, and analyzed mouse fate mapping experiments. ZC performed mouse bioinformatic analyses and assisted in the preparation of related figures. SN, AKH and EGP provided expertise for data analysis and interpretation. AAG provided general support. All authors participated in the writing of the manuscript.

## Competing interests

The authors declare no competing interests.

## Data and materials availability

Materials and additional details will be provided upon request, under material transfer agreements with MSKCC.

## Notes

### Competing Interest Statement

The authors have declared no competing interest.

