## Supplementary material for "Identification of a mesodermal progenitor for the pro-definitive angio-hematopoietic lineage": methods and extended data

**Lazarov et al.,**
**Methods and Extended Figures**

**The PDF file includes:**

Methods
Extended Data Figures 1 to 8
Supplementary References

### Methods

#### Mice

*$Cxcr4^{Egfp/+}$*[1], *$Cxcr4^{CreERT2}$*[2], and *$Myb^{-/-}$*[3] were previously described. *$Csf1r^{–/–}$* mice[4] on FVB/NJ background were provided by E.Richard Stanley. B6;129S6-*$Gt(ROSA)^{26Sortm14(CAG-tdTomato)Hze}$*/J (*$Rosa26^{LSL-tdTomato}$*, JAX Stock No: 007908)[5,6] and *$Gt(ROSA)26Sor^{tm4(ACTB-tdTomato,-EGFP)Luo}$/J (Rosa26$^{mT/mG}$*, JAX Stock No: 007576)[7] were purchased from Jackson laboratories. All mice were maintained under specific-pathogen-free conditions and experiments were performed in accordance with animal license issued by the Institutional Review Board (IACUC 15-04-006) at Memorial Sloan Kettering Cancer Center (MSKCC)

List of mouse strains and genotyping protocols:

| Strain | Genotype Primers | Sequence (5'-3') | Expected band | Dena. | Ann. | Elon. | Cyc. | Final Elon. |
|---|---|---|---|---|---|---|---|---|
| *$CXCR4^{GFP/-}$* | Cxcr4-F | AGC AGT GAA ACC TCT GAG GCG TT | WT allele: no band | 94°C 30sec | 60°C 30sec | 72°C 30sec | 35 | 72°C 7 min |
| | Cxcr4-R | CGG CGA GCT GCA CGC TGC CGT CCT C | Mut allele: 600bp | | | | | |
| *$CXCR4^{EGFP-CreERT2}$* | Cxcr4CreERT2_U | AGT GAA ACC TCT GAG GCG TTT GGT | WT allele:300bp | 95°C 30 sec | 60°C 30sec | 72°C 30sec | 35 | 72°C 5 min |
| | Cxcr4CreERT2_L1 | TAG AGC CTG TTT TGC ACG TTC ACC | Mut allele:500bp | | | | | |
| | Cxcr4CreERT2_L2 | TCT GAA CCC GTC CCA CTC AAC TTA | | | | | | |
| *$Rosa26^{LSL-tdTomato}$* | WT allele | AAGGGAGCTGCAGTGGAGTA | Wild allele:297 bp | 95°C 30sec | 60°C 30sec | 72°C 30sec | 40 | 72°C 5 min |
| | WT allele | CCGAAAATCTGTGGGAAGTC | | | | | | |
| | Mutant allele | GGCATTAAAGCAGCGTATCC | Mut allele: 196 bp | | | | | |
| | Mutant allele | CTGTTCCTGTACGGCATGG | | | | | | |
| *$Rosa26^{LSL-mT/mG}$* | WT R | GGA GCG GGA GAA ATG GAT ATG | WT allele: 374 bp | 94°C 40sec | 65°C 60sec | 72°C 60sec | 28 | 72°C 5 min |
| | WT | GGC TTA AAG GCT AAC CTG ATG TG | | | | | | |
| | Mutant R | AAT CCA TCT TGT TCA ATG GCC GAT C | Mut allele: 500 bp | | | | | |
| | Mutant | CCG GAT TGA TGG TAG TGG TC | Heterozygote: 374bp &500bp | | | | | |
| *$Csf1r^{-/-}$* | P1 WT | TCTCCTGGGATGGGAAACGATCCCAAAGGC | WT allele: 536bp | 94°C 30sec | 60°C 30sec | 72°C 30sec | 35 | 72°C 7 min |
| | P2 WT | GATTCAGGGTCCAAGGTCCAGATGGGAGAG | | | | | | |
| | P3 Mut | GCCAGCCACGATAGCCGCGCTGCCTCGTC | Mutant allele: 1.6Kb | | | | | |
| | P4 Mut | CTT CCT GGC CCT CAA CCA CTG TCA | | | | | | |
| | WT R allele | TGC TGT CCT TCA TGT CGC CG | | | | | | |
| *$Myb^{-/-}$* | c-myb null allele | CCA TGC GTC GCA AGG TGG AAC | Mut allele: 295 bp | 94°C 30sec | 60°C 30sec | 72°C 30sec | 35 | 72°C 7 min |
| | c-myb null allele R | TGG CCG CTT TTC TGG ATT CAT C | | | | | | |

#### Mouse fate mapping studies

Mice were housed under a 12-hour light/dark cycle. Natural mattings were set-up between males and 6–8-week-old virgin females, with noon of the day of vaginal plug considered embryonic day (E) 0.5. For the analysis of *Cxcr4* expression, *$Cxcr4^{Egfp}$* male reporter mouse on C1D background[1] were mated to C1D female mice. For *Cxcr4* expression fate mapping studies, *$Cxcr4^{CreERT2}$*[2] male on C57BL/6J background were crossed with *$Rosa26^{LSL-tdTomato}$*[6] or *$Rosa26^{mTmG}$*[7] reporter female mice. To induce Cre-mediate recombination, *$Rosa26^{LSL-tdTomato}$ or $Rosa26^{mT/mG}$* pregnant mothers mated to *$Cxcr4^{CreERT2}$* received a single i.p. (intra peritoneal) injection of 4-hydroxytamoxifen (OH-TAM) (20µg per gram of body weight of the pregnant mother) supplemented with progesterone (37.5µg per gram body weight), at post coitum timepoints ranging from E5.75 to E10.5, as previously described[2]. Following embryonic Cre recombination, *$Cxcr4^{CreERT2}$; $Rosa26^{LSL-tdTomato}$* mice were sacrificed at time points ranging from E7.5 to E15.5 or after birth at 4 weeks of age for the analysis of tdTomato+ cells in different tissues by flow cytometry. *$Cxcr4^{CreERT2}$; $Rosa26^{mTmG}$* mice were sacrificed at E10.5 and at 4 weeks of age and tissues were analyzed by immunofluorescence imaging.

#### Mouse immunofluorescence imaging

Analysis of Cxcr4 expression during embryonic development was performed using Wholemount Immunofluorescence microscopy of *Cxcr4*$^{gfp}$ reporter embryos at embryonic days E7.25 and E8.25. For E7.25 embryos the Reichert's membrane was carefully removed prior to fixation. Whole embryos were fixed for 30 min with 4% PFA at room temperature and then incubated with PBS 1% BSA, 0.1% Triton X-100 (Sigma) and 10% Goat serum (Sigma) for 1 hour. The embryos were stained with chicken anti-GFP (1:1000 Aves Labs), Rat anti-CD309(KDR) (1:100 eBioscience 14-5821-82), Rabbit anti-Brachyury (1:500 abcam, ab209665), Armenian Hamster anti-CD31 (1:100 Thermo Fisher, MA3105) overnight at 4ºC in PBS 1% BSA, 0.1% Triton X-100 (Sigma). After 3 washes in PBS 1% BSA, 0.1% Triton X-100 (Sigma) and 10% Goat serum the embryos were then incubated with the following secondary antibodies for 3 hours at room temperature: Goat anti-Rabbit Alexa fluor 405, Goat anti-Chicken Alexa Fluor 488 (1:500; Invitrogen, A11039), Goat anti-Armenian hamster Alexa Fluor 647 (1:200 Biolegend, 405510), Goat anti-Rat Alexa Fluor 568 (1:200 Thermo Fisher Scientific, A11077). Embryos were washed 3 times in PBS 0.1% Triton X-100 (Sigma). Prior to imaging E7.25 embryos were carefully orientated to image the posterior side while E8.25 embryos were orientated to image the lateral side. Embryos were immobilized in custom agarose molds in 35mm µ-Dish (Ibidi). Imaging was performed using LSM880 confocal microscope with Plan-Apochromat 20x/0.8 objective (Zeiss). Analysis of GFP+ KDR+ double positive and GFP+ CD31+ double positive cells was performed by generating 3D surface rendering for each channel and using Surface-surface colocalization XTension in Imaris software (Bitplane). All image analysis will be performed using Imaris software (Bitplane).

Fate-mapping of Cxcr4 expressing cells during embryonic development was performed using Immunofluorescence microscopy of *Cxcr4*$^{CreERT2}$*; Rosa26*$^{mT/mG}$ embryos at E10.5 or at 4 weeks of age after OH-TAM treatments at E6.5 or E8.25 (see mouse fate mapping studies section). After dissecting the placenta, the yolk sac was separated from the embryo proper before fixation. Embryos were fixed overnight with 4% PFA whereas yolk sacs were fixed for 30min at room temperature. Yolk-sac were preserved in PBS for wholemount imaging while embryos were processed for Cryosection. Embryos were washed in PBS and dehydrated overnight in 30% Sucrose in PBS. Embryos were embedded in FSC22 Frozen section Media (Leica). Embryo samples were frozen at -80˚C and sectioned using a Leica CM3050S cryostat. Cryoblocks were cut at a thickness of 16µm, placed on SuperFrost Plus slides (Fisherbrand). Embryo sections and whole yolk sacs were blocked with PBS 1% BSA and 0.3% Triton X-100(Sigma) for 1 hour at room temperature. Samples were incubated with Chicken anti-GFP (1:200 Aves Labs), and Rat anti-CD31 (1:100 BD bioscience Cat:550274) overnight at 4ºC. After washing with 1% BSA and 0.3% Triton X-100 PBS, secondary antibody staining was performed using Goat anti-chicken Alexa Fluor 488 (1:200; Invitrogen, A11039) and Goat anti-Rat Alexa fluor 647 (1:200 Invitrogen, A21247) incubation for 2 hours at room temperature. After nuclei staining with Hoechst 33342 (1:1000, ThermoScientific) for 10 mins, the samples were washed 3 times and mounted using ProLong Gold antifade reagent (Invitrogen, Ref P36930). Brain and Liver tissue from 4 weeks old *Cxcr4*$^{CreERT2}$*; Rosa26*$^{mT/mG}$ mice that received OH-TAM treatments at E6.5 were analyzed by whole mount immunofluorescence. Whole Brain and Liver were fixed for 30 min at room temperature. Brain and Liver wholemount were were blocked with PBS 1% BSA and 0.3% Triton X-100(Sigma) for 1 hour at room temperature. Samples were incubated with Chicken anti-GFP (1:200 Aves Labs), Rat anti-CD11b for the brain (1:100 BD bioscience Cat:550282) or Rabbit anti-F4/80(1:200 Cell Signaling Cat:30325) for the liver overnight at 4ºC. After washing with 1% BSA and 0.3% Triton X-100 PBS, secondary antibody staining was performed using Goat anti-chicken Alexa Fluor 488 (1:200; ThermoFisher Scientific, A11039), Goat anti-Rat Alexa fluor 647 (1:200 ThermoFisher Scientific, A21247) or Goat anti-Rabbit Alexa Fluor 647 (1:200 ThermoFisher Scientific A21245) incubation for 2 hours at room temperature. After nuclei staining with Hoechst 33342 (1:1000, ThermoScientific) for 10 mins, the samples were washed 3 times and mounted using ProLong Gold antifade reagent (Invitrogen, Ref P36930). Embryo sections, yolk sacs and adult tissues whole mounts were imaged using LSM880 confocal microscope with oil immersion 40x 1.3 N.A. objective and 1µm Z-stack intervals. Image analysis was performed using Imaris software (Bitplane).

Mouse flow cytometry

To prepare samples for flow cytometry analysis, whole embryos or individual embryonic tissues were dissected, minced into small pieces, and incubated in digestion buffer containing 1X PBS, Collagenase D (1mg/ml, Sigma), DNAse (0.2mg/ml, Sigma) and 3% heat-inactivated fetal bovine serum (FBS, Invitrogen) for 15min at 37°C. Digested suspensions were further dissociated by passing them through a 100µm cell strainer (BD). Single cell suspensions were centrifuged (400g, 5minutes, 4°C), and resuspended in 50µl of blocking buffer containing 1X PBS, 0.5%BSA, 2mM EDTA, anti-mouse CD16/32 (1:100), for 15min at 4°C. Samples were then stained with indicated antibodies for 30min at 4°C.

Once processed, all samples were resuspended in FACS buffer (PBS, 0.5% BSA, and 2 mM EDTA) containing anti-mouse CD16/32 (FcRIII/II, Biolegend 101302) and anti-mouse CD16.2 (Biolegend 149502) for 10 min and stained with antibody mixes for 30 min on ice. After two washes with FACS buffer, samples were incubated with 2 µM Hoechst 33342 (Thermo Scientific) just prior to analysis on a LSR Fortessa X-20 (BD Biosciences) or sorting on an ARIA III (BD Biosciences). All flow cytometry data was analyzed by Treestar FlowJo V9.9 or V10.10.

Mouse Flow Cytometry Antibodies

| Antibody | Dilution | Clone | Conjugate | Company |
|---|---|---|---|---|
| FLK-1 (KDR) | 1:100 | Avas 12a1 | BV605 | BD Pharmingen |
| CD31 (PECAM-1) | 1:100 | MEC13.3 | BV786 | BD Pharmingen |
| CD115 | 1:100 | T38-320 | BV650 | BD Pharmingen |
| CD117 (c-KIT) | 1:100 | 2B8 | APC Cy7 | Biolegend |
| CD11b | 1:200 | M1/70 | PE Cy7 | BD Pharmingen |
| CD150 | 1:200 | TC15-12F12.2 | PE Cy7 | Biolegend |
| CD16/32 | 1:100 | 2.4G2 | Uncoupled | BD Pharmingen |
| CD41 | 1:100 | MWReg30 | AF700 | Biolegend |
| CD19 | 1:500 | 1D3 | BV711 | BD Pharmingen |
| CD3 | 1:100 | 145-2C11 | BV711 or APC Cy7 | ebioscience |
| CD335 (NKp46) | 1:100 | 29A1.4 | BV711 | ebioscience |
| Cd369 (Clec7A, Dectin-1) | 1:100 | bg1fpj | APC | ebioscience |
| CD45 | 1:100 | 30-F100 | AF780 | ebioscience |
| CD48 | 1:100 | HM48-1 | APC | Biolegend |
| CD93 | 1:100 | AA4.1 | APC | Biolegend |
| Gr1 | 1:200 | RB6-8C5 | BV711 | BD Pharmingen |
| Ly-76 | 1:100 | Ter119 | BV711 or AF700 | BD Pharmingen |
| Ly6C | 1:100 | HK1.4 | BV786 | Biolegend |
| Ly6G | 1:200 | 1A8 | BV510 | BD Pharmingen |
| NKp46 | 1:100 | 29A1.4 | AF647 | BD Pharmingen |
| Sca1 | 1:200 | D7 | BV510 | BD Pharmingen |
| F4/80 | 1:200 | 2M8 | eF450 or APC or BV605 | ebioscience |
| Tim-4 | 1:200 | RMT4-54 | AF647 | BD Bioscience |
| CD64 | 1:200 | 10.1 | APC | BD Bioscience |

Analysis of public single-cell RNA-seq datasets

Mouse scRNA-seq data from Pijuan-Sala et. al., Nature 2019 [8], including scRNAseq read counts, cell type annotations, and time point information was obtained from the following GitHub repository: https://github.com/MarioniLab/EmbryoTimecourse2018. Cells were subset to include only the following populations: epiblast, caudal epiblast, primitive streak, nascent mesoderm, mixed mesoderm, hematoendothelial progenitors, endothelium, blood progenitors, and erythroid cells. We defined cells as being CXCR4 or KDR positive if their expression was at or above the 90th percentile for expression CXCR4. Differential gene expression was performed between defined groups (E7 Kdr+Cxcr4+ vs E7 Kdr+Cxcr4-) using the scanpy function scanpy.tl.rank_genes_groups; statistical testing was performed using the Wilcoxon rank sum test (option method = 'wilcoxon') and multiple hypothesis testing correction was performed by applying the Benjamini-Hochberg correction (option corr_method = 'benjamini-hochberg'). For visualization, we selected the top five thousand highly variable genes (scanpy.pp.highly_variable_genes) and performed library size normalization and log transformation with a pseudocount of 1E-5. We then ran UMAP (scanpy.pl.umap) on the top thirty principal components after running principal components analysis (PCA). For visualization only (and not for differential expression analysis), log library size-normalized gene counts were imputed by running MAGIC (scanpy.external.tl.magic), using the top 30 principle components as the input embedding (n_pca = 30) and five nearest neighbors for graph construction (knn = 5). Human scRNAseq data from Human Embryo, Carnegie stage 7 from Tyser et. al., Nature 2021[9] was analyzed using the tools provided by the authors at: www.human-gastrula.net.

Derivation and quality control of human induced pluripotent stem cell (hiPSC) lines

HiPSC lines were derived from frozen peripheral blood mononuclear cells (PBMCs) of three independent donors. Written informed consent was obtained according to the Helsinki convention (Ethics approval: 11/LO/1433). The study was approved by the Institutional Review Board of St Thomas'Hospital; Guy's hospital; the King's College London University and the Memorial Sloan Kettering Cancer Center. hiPSC were derived according to published protocols[10] using Sendai viral vectors (ThermoFisher Scientific; A16517). Newly derived iPSC clones are maintained in culture for 10 passages (2-3 months) to remove any traces of Sendai viral particles and ensure the cells remain stable during prolonged culturing period. Over 90% of iPSCs in the derived lines expressed high levels of the pluripotency markers NANOG and OCT4 by flow cytometry. Karyotyping analysis showed normal karyotype (46, XX). iPSC lines tested negative for the presence of mycoplasma using MycoAlert Plus kit (Lonza). iPSC clones that passed all quality controls were frozen down and used for downstream experiments.

Generation of mutant iPSC lines

CRIPSR-CAS9 technology was used to generate mutant RUNX1, NOTCH1, MYB, and CSF1R hiPSCs lines. CRISPR sgRNA targeting each gene were designed using the algorithm available at https://www.benchling.com/crispr/. The target sequences were cloned into a pX330-U6-Chimeric_BB-CBh-hSpCas9 vector (Addgene plasmid #42230) to make specific gene targeting constructs. To introduce mutations in the parental line (C12 WT iPSC Line), iPSC single cell suspensions were electroporated ($1x10^6$ cells per reaction) with 4μg sgRNA-construct plasmid using Human Stem Cell NucleofectorTM solution (Lonza) following manufacturer's instructions. The electroporated cells were seeded and cultured for 4 days in feeder free conditions using Essential 8 medium in accordance to manufacturer's protocol (Thermo Fisher, A1517001). The cells were then dissociated by Accutase (Fisher Scientific, A1110501) and re-plated at a low density (4 cells per well in 96-well plates) to get single-cell clones. 10 days later, individual colonies were picked, expanded and analyzed by PCR and DNA sequencing. The sgRNA target sequences, PCR primers, and Sequencing primers are listed below.

iPSC Line sgRNA target sequences and PCR and sequencing primers

| Gene | sgRNA Target | PCR-Forward Primer | PCR-Reverse Primer | Sequencing Primer |
|---|---|---|---|---|
| RUNX1 | CACTTCGACCGACAAACCTG | AGATGGCACTCTGGTCACTG | TCACAATTCCTACGTTGCATGT | TGTGATGGCTGGCAATGATG |
| NOTCH1 | CTTCACACTTCCCGCCATTC | GTTGACGCCCCTTTGGTTTC | AGCTCCACACGCAGCATAAT | GACGCCCCTTTGGTTTCATGG |
| MYB | AAGTCTGGAAAGCGTCACTT | TTGGCCGTGTCTGTGGATAC | GAGGTGATGCCTTCATGGGT | GGATACCCATAACAGCAGAACCA |
| CSF1R | AACGGTGACCTTGCGATGTG | CCAAGCTACCACCCATCTCA | CTCTTCCAGGTCCTTGCTCA | ACTTCTGTCTGCTGTCCCTC |

Culture and differentiation of hiPSCs

HiPSCs were maintained on irradiated CF1 mouse embryonic fibroblasts (MEFs, ThermoFisher Scientific; A34181) in Embryonic Stem Cell (ESC) medium supplemented with 10 ng/ml basic fibroblast growth factor (bFGF, Peprotech; 100-18B). Media was changed every other day. Passaging was performed every 7 days at a dilution ratio of 1/4 to 1/10, depending on colony density and size. During passaging, iPSCs were detached as clusters by a 13 min incubation at 37°C with collagenase type IV (250 UI/ml final concentration) (ThermoFisher Scientific; 17104019) and were pelleted at room temperature by a 5-minute centrifugation at 150G. The iPSC clusters were resuspended in ESC medium supplemented with 10ng/ml bFGF (Peprotech; 100-18B) and plated on NUNC plates containing pre-plated (2 days in advance) 12,500 to 16,000 MEFs per $cm^2$.

HiPSC hematopoietic differentiation was adopted from a previously published protocol[11] and modified as follows. At day 0 of the differentiation, expanded hiPSCs were detached as described above and transferred (from 150mm plate, to 4wells) for cultivation in 6 well low adhesion plates in ESC media supplemented with 10µM ROCK Inhibitor (Sigma; Y0503). The plates were kept on an orbital shaker at 100rpm for 6 days to allow for a spontaneous formation of embryoid bodies (EB) with hematopoietic potential. At day 6 of the differentiation, 200-500µm cystic EBs were picked under a dissecting microscope and transferred onto adherent tissue culture plates (~2.5 EBs/cm2) for cultivation in Hematopoietic Differentiation (HD) medium from day 6 up to day 18 from the start of differentiation, supplemented with hIL-3 (Peprotech; 200-03) (25ng/ml) and hM-CSF (Peprotech, 300-25) (50ng/ml). All cells were cultured in incubators at 37°C in a humidified atmosphere of 5% $CO_2$.

hiPSC culture and differentiation media components

| Medium Name | Reagent | Supplier; Catalog Number | Final Conc. |
|---|---|---|---|
| Embryonic Stem Cell (ESC) Medium | Knock-Out Dulbecco's Modified Eagle Medium | Thermo Fisher Scientific 10829-018 | / |
| | KO serum replacement | Thermo Fisher Scientific 10828-028 | 20% |
| | L-glutamine | Thermo Fisher Scientific; 25030-024 | 2mM |
| | Nonessential Amino Acids | Thermo Fisher Scientific; 11140-035 | 1% |
| | Penicillin/Streptomycin | Thermo Fisher Scientific, 15140163 | 1% |
| | B-mercaptoethanol | Thermo Fisher Scientific; 31350-010 | 0.2% |
| Hematopoietic Differentiation (HD) Medium | APEL 2 | Stem Cell Tech; 05270 | / |
| | Protein Free Hybridoma | Thermo Fisher Scientific; 12040077 | 5% |
| | Penicillin/Streptomycin | Thermo Fisher Scientific, 15140163 | 1% |

Digestion of EB/iPSC clusters for flow cytometry analysis and cell sorting

To digest EBs to a single cell suspension for analysis and cell sorting by flow cytometry, EB/iPSC clusters and surrounding media were collected from day 0 to day 18 of the differentiation cultures, washed with room temperature PBS, and pelleted by a 5-minute room temperature centrifugation at 800G. The pellet was resuspended in TrypLE at 37ºC (Thermo-Fischer 12605-010) and incubated for 10 minutes. After 10 minutes the suspension was pipetted with a 1ml pipette 30 times, left for 1 minute to allow for undigested EB clusters to sediment, and the top part of the liquid containing single cells in suspension was transferred through a 100µm mesh filter into a tube containing room temperature IMDM media (Thermo Scientific, 12440053) with 10% FBS (EMD Milipore, TMS-013-B). This cycle was repeated 4 times until the EBs were fully digested. The single cell suspension was centrifuged (400G, 5minutes, 4C), and resuspended in FACS buffer (PBS, 0.5%BSA, 2mM EDTA) at a density of $<1x10^7$ cells/ml.

The cell suspension was incubated with Fc block (BD 564219) at a 1/20 dilution, at room temperature for 10 minutes. The cells were then washed in FACS buffer, resuspended to a density of $<1x10^7$ cells/ml, and stained for 30 minutes at 4ºC with antibodies listed below. The cells were washed and resuspend in FACS buffer to a density of $1x10^6$ cell/ml. DAPI (1ng/ml final) was added for dead cell exclusion immediately prior to flow cytometry acquisition and cell sorting done on BD Aria III. Collection media for sorted cells varied depending on the downstream procedures and has therefore been specified within the description of those experimental protocols. Full gating strategy for cell sorting of pop1-10 is shown in Extended Data Fig. 5. The total number of cells per well was determined using a cell counter (GUAVA easyCyte HT).

Flow Cytometry Anti-Human Antibodies:

| Antibody | Dilution | Clone | Manufacturer | Reference |
|---|---|---|---|---|
| CD235a BV510 | 1:100 | HIR2 | BD | 740174 |
| CD45 BV650 | 1:50 | HI30 | BD | 563717 |
| CD11b BV711 | 1:100 | ICRF44 | BioLegend | 301344 |
| CD11b PE-Cy7 | 1:100 | ICRF44 | BioLegend | 301322 |
| CD144 (VE-Cadh) BV786 | 1:100 | 55-7H | BD | 565672 |
| CD206 (MRC1) AF488 | 1:100 | 19.2 | eBioscience | 53-2069-42 |
| CD66b PE | 1:100 | G10F5 | BioLegend | 305106 |
| CD34 PerCp-Cy5.5 | 1:100 | 8G12 | BD | 347203 |
| CD309 (KDR) PE-Cy7 | 1:250 | 7D4-6 | BioLegend | 359912 |
| CD184 (CXCR4) APC | 1:400 | 12G5 | BioLegend | 306509 |
| CD41 AF700 | 1:100 | HIP8 | BioLegend | 303727 |
| CD14 – APC-Cy7 | 1:100 | M5E2 | BioLegend | 301820 |
| CD43 PerCp-Cy5.5 | 1:100 | CD43-10G7 | BioLegend | 343205 |
| CD31 PerCp-Cy5.5 | 1:100 | WM59 | BioLegend | 303131 |
| CD19 APC | 1:100 | HIB19 | BioLegend | 302211 |
| CD3 PerCp-Cy5.5 | 1:100 | UCHT1 | BD | 560835 |
| NKp46 PerCp-Cy5.5 | 1:100 | 9E2 | BioLegend | 331919 |

Summary of gating strategy for sorted cells:

| Population | Gates |
|---|---|
| 1 | CD206-, CD14-, CD45-, CD144-, CD235a-, CD41-, CXCR4-, KDR+ |
| 2 | CD206-, CD14-, CD45-, CD144-, CD235a-, CD41-, CXCR4+, KDR+ |
| 3 | CD206-, CD14-, CD45-, CD144-, CD235a+, CD41-, |
| 4a | CD206-, CD14-, CD45-, CD144+, CD235a-, CXCR4+ |
| 4b | CD206-, CD14-, CD45-, CD144+, CD235a-, CXCR4- |
| 4c | CD206-, CD14-, CD45-, CD144+, CD235a+, CD41- |
| 4d | CD206-, CD14-, CD45-, CD144+, CD235a+, CD41+ |
| 5 | CD206-, CD14-, CD45+, CD144-, CD41+ |
| 6 | CD206-, CD14-, CD45+, CD144-, CD235a-, CD41-, CD11b-, CD66b- |
| 7 | CD206-, CD14-, CD45+, CD144-, CD235a+, CD41- |
| 8 | CD206-, CD14-, CD45+, CD144-, CD235a-, CD41-, CD11b+, CD66b- |
| 9 | CD206+, CD14+, CD45+ |
| 10 | CD206-, CD14-, CD45+, CD144-, CD235a-, CD41-, CD66b+ |

Flow cytometry tSNE analysis

HiPSC flow cytometry data were analyzed with FlowJo 9.9. For t-SNE analysis of the differentiation cultures, $6x10^4$ single live cells per timepoint (determined by gating on FSC-A/SSC-A, FSC-A/FSC-W, and DAPI negative) from day 0-6, or day 6-18 were concatenated. Within the concatenated files, cells negative for all markers as determined by Fluorescence Minus One (FMO) controls were excluded by a Boolean gate and t-SNE analysis was conducted on the remaining cells (positive for at least one marker). t-SNE analysis was conducted in FlowJo 9.9 with parameters set to 600 iterations, perplexity of 20 and theta of 0.5. The t-SNE analysis for day 0-6 timepoints included the following flow cytometry fluorescence channels/antibodies: FSC-A, SSC-A, KDR, CXCR4, CD235a, CD144, CD41, CD45, CD11b, CD14, CD206, CD66b. The t-SNE analysis for day 6-18 timepoints included all of the fluorescence channels/antibodies listed above except KDR and CXCR4 as these antibodies were not included in the flow cytometry panel for those particular experiments. Clusters were annotated manually based on t-SNE separation and antibody expression patterns.

OP9 co-culture cell differentiation potential assays

Protocols for coculture of sorted populations with OP9 stromal cells to evaluate differentiation potential were adopted from previously published studies[12,13] and modified as follows. OP9 cells were purchased from ATCC (CRL-2749) and cultured in accordance with ATCC recommendations in OP9 Maintenance media (See “OP9 coculture cell differentiation potential assay media composition” list below). OP9 cells at 90% confluency were treated with 10µg/ml of Mitomycin C (Cayman Chemical Company, 11435) for 3 hours. After treatment, the cells were washed 2 times with PBS to remove any traces of Mitomycin C, detached using TrypLE for 3 minutes, pelleted by a 5-minute 300G centrifugation at room temperature, and frozen down at $1x10^6$ cells per vial in 500µl of Stem-CellBanker solution (Thermo Fisher Scientific, NC0261714). Mitomycin C treated OP9 cells were thawed, counted, and seeded at a density of $1x10^5$ cells per $cm^2$ on tissue culture plates in OP9 Maintenance Media, one day before FACS-isolated cells were added.

Day 3 or Day 15 iPSC differentiation cultures were processed for flow cytometry as described above (see Digestion of EB/iPSC clusters for flow cytometry analysis and cell sorting section) and population 1-10 were sorted based on their expression phenotypes described above, into 1.5ml Eppendorf tubes (precoated for 2hrs with PBS 20% FBS), containing 500µl of collection media (PBS, 2% FBS, 25mM HEPES buffer) at room temperature. The sorted populations were pelleted by a 5-minute, 400G centrifugation at room temperature and resuspended in OP9 Differentiation media.

To evaluate the differentiation potential of FACS-isolated cells by flow cytometry, 1000-2000 live cells of pop1-10 were added to 12 well tissue culture plates (MIDSCI, TP92412) containing Mitomycin C treated OP9 cells. The co-culture of OP9 cells and FACS-isolated cells was maintained in OP9 Differentiation media for 1, 5, 7, or 14 days. Media was changed on day 7. For flow analysis of the co-cultures, the cells were detached into a single-cell suspension using a 5-minute incubation with TrypLE. The cell suspension was processed for flow cytometry as described above (see the second paragraph of the "Digestion of EB/iPSC clusters for flow cytometry analysis and cell sorting" section). The antibody staining panel included KDR, CXCR4, CD235a, CD144, CD41, CD45, CD11b, CD206, CD66b, as well as an anti-mouse CD29 antibody (1:100, Biolegend, 102228) to exclude OP9 cells from the analysis. The samples were acquired in their entirety to count the absolute number of cells per well.

To evaluate the clonal differentiation potential of FACS isolated population 1, 2, 4a, 4b, 4c, or 4d, each cell population was plated at an estimated density of one live cell per well in Bio-one 384 well µClear imaging plates (Fisher Scientific, 07-000-053) containing pre-plated OP9 cells. The co-culture of OP9 cells and FACS-isolated cells was maintained in OP9 Differentiation media for 5 days. Each well was then fixed for 10 min in 4% PFA. The samples were blocked in PBS 1% BSA and 0.3% Triton X-100(Sigma) 10% Goat serum (Sigma) for 1 hour at room temperature. Samples were incubated with Anti-human Mouse CD144 (1:100, BD Bioscience, 555661), anti-human Rat CD235a (1:50, Bio-rad, MCA506G), anti-human Rabbit CD14 (1:100, Abcam ab221678) overnight at 4ºC. After washing with 1% BSA, 0.3% Triton X-100 PBS and 10% Goat serum (Sigma), secondary antibody staining was performed using Goat anti-Mouse Alexa fluor 488 (1:200, Thermo Fisher Scientific, A11029), Goat anti-Rat Alexa Fluor 568 (1:200, Thermo Fisher Scientific, A11077), Goat anti-Rabbit Alexa Fluor 647 (1:200, Thermo Fisher Scientific A32733) for 3 hours at room temperature. The samples were washed 3 times in PBS 0.3% Triton X-100 (Sigma). PBS was added to the stained samples, and the entire wells were imaged. Imaging was performed using LSM880 confocal microscope with Plan-Apochromat 20x/0.8 objective (Zeiss) and 2.5 µm Z-stack intervals. Each well was then analyzed for the presence or absence of pop 3 ($CD235a^+$, $CD144^-$, $CD14^-$), pop 4 ($CD144^+$ vessels), or pop 10 ($CD14^+$ $CD144^-$ $CD235a^-$) cells using Imaris software (Bitplane).

OP9 coculture cell differentiation potential assay media composition

| Media | Reagent | Supplier; Catalog Number | Final Concentration |
|---|---|---|---|
| OP9 Maintenance Media | Alpha Minimum Essential Medium without ribonucleosides and deoxyribonucleosides | Thermo Scientific, 12561056 | / |
| | FBS (not heat treated) | EMD Milipore, TMS-013-B | 20% |
| | Penicillin/Streptomycin | Thermo Fisher Scientific; 15140163 | 1% |
| OP9 Differentiation Media | OP9 Maintenance Media | See rows above | / |
| | rhVEGF | R&D Systems, 293-VE-010 | 5ng/ml |
| | rhSCF | R&D Systems, 255-SC-010 | 50ng/ml |

| | | | |
|---|---|---|---|
| | rhTPO | PeproTech, 300-18 | 30ng/ml |
| | rhFlt3 | Stem Cell Technologies, 78009.1 | 10ng/ml |
| | rhIL-11 | PeproTech, 200-11 | 5ng/ml |
| | rhBMP-4 | R&D Systems, 314-BP-010 | 10ng/ml |
| | rhIL-3 | PeproTech, 200-03 | 10ng/ml |
| | rhIL-6 | R&D Systems, 206-IL-010 | 20ng/ml |
| | rhEPO | PeproTech, 100-64 | 2U/ml |
| | Human Holo-Transferrin | Sigma-Aldrich, T0665-50MG | 150µg/ml |

Cells differentiation potential assays on MethoCult

Day 3 or Day 15 iPSC differentiation cultures were processed for flow cytometry as described above (see "Digestion of EB/iPSC clusters for flow cytometry analysis and cell sorting" section) and population 1-10 were sorted based on their expression phenotypes described in Figure 3, into 1.5ml Eppendorf tubes (precoated for 2hrs with PBS 20% FBS), containing 500µl of Iscove's MDM with 2% FBS (Stem Cell Technologies, 07700) at room temperature. The sorted populations were pelleted by a 5-minute, 400G centrifugation at room temperature and resuspended and plated according to manufacturer's protocol in serum-free MethoCult SF H4636 methylcellulose (Stem Cell Technologies, 04636). After 14 days of culture, the contents of the plates were collected, washed with PBS, and processed for flow cytometry as described above, in the second paragraph of the "Digestion of EB/iPSC clusters for flow cytometry analysis and cell sorting" section. The samples were acquired in their entirety in order to count the absolute number of cells per replicate/plate.

Whole mount immunofluorescence imaging of hiPSCs derived EBs

HiPSCs derived EBs, generated as described above (see "Culture and differentiation of hiPSCs" section), were imaged at day 3 and day 12 of the differentiation culture. iPSC/EB clusters were collected (for day 3 iPSC/EB clusters), or directly fixed on plates (for day 12 EB cultures) after washing with PBS, using 4% PFA for 10min at room temperature. The samples were blocked in PBS 1% BSA and 0.3% Triton X-100(Sigma) 10% Goat serum (Sigma) for 1 hour at room temperature.

Day 3 cultures were stained overnight at 4ºC with Rabbit anti-human CXCR4 (1:200, Abacm, ab 124824), Mouse Anti-human CD144 (1:100, BD, Bioscience 555661), and Rat anti-human CD235a (1:50, Bio-rad, MCA506G), or with Rat anti-human CXCR4 (1:200 BD Biosciences 551413), Mouse Anti-human CD144 (1:100, BD, Bioscience, 555661), and Rabbit anti-human VEGF receptor 2 20(KDR, 1:100, Abcam ab11939) antibody panel combinations. After washing with 1% BSA, 0.3% Triton X-100 PBS and 10% Goat serum (Sigma), secondary antibody staining was performed using Goat anti-Mouse Alexa fluor 488 (1:200, Thermo Fisher Scientific, A11029), Goat anti-Rat Alexa Fluor 568 (1:200, Thermo Fisher Scientific, A11077), Goat anti-Rabbit Alexa Fluor 647 (1:200, Thermo Fisher Scientific, A32733) for 3 hours at room temperature. After nuclei staining with Hoechst 33342 (1:1000, Thermo Fisher Scientific) for 10 mins, the samples were washed 3 times in PBS 0.3% Triton X-100(Sigma). Day 3 whole iPSC/EB clusters were mounted on imaging slides using 0.12mm depth Secure seal spacers (Electron Microscopy Sciences, Cat#70327-20s) in ProLong Gold antifade reagent (Invitrogen, Ref P36930).

Day 12 cultures were stained overnight at 4ºC with Anti-human Mouse CD144 (1:100), anti-human Rat CD235a (1:50), anti-human Rabbit Integrin α2b (CD41, 1:100, Cell signaling 13807S). After washing with 1% BSA, 0.3% Triton X-100 PBS and 10% Goat serum (Sigma), secondary antibody staining was performed using Goat anti-Mouse Alexa fluor 488 (1:200, Thermo Fisher Scientific, A11029), Goat anti-Rat Alexa Fluor 568 (1:200, Thermo Fisher Scientific, A11077), Goat anti-Rabbit Alexa Fluor 647 (1:200, Thermo Fisher Scientific, A32733) for 3 hours at room temperature. After nuclei staining with Hoechst 33342 (1:1000, Thermo Fisher Scientific) for 10 mins, the samples were washed 3 times in PBS 0.3% Triton X-100 (Sigma) and filled with PBS to prevent drying. Imaging was performed using LSM880 confocal microscope with Plan-Apochromat 20x/0.8 objective (Zeiss) and 2.5 µm Z-stack intervals.

Cytology (Cytospin preparation and staining)

Day 3 or Day 15 iPSC differentiation cultures were processed for flow cytometry as described above and 1000-5000cells of population 1-10 were sorted based on their expression phenotypes described in Figure 3, into 1.5ml Eppendorf tubes (precoated for 2hrs with PBS 20% FBS), containing 200µl of FBS at room temperature. The collected cells were loaded into Cytofunnels (Fisher Scientific, BMP-CYTO-DB25) attached to Super-frost slides (Thermo Scientific, 12-550-15), and were spun down with Cytospin 3 (Thermo Shandon) centrifuge at 800rpm for 10min (medium acceleration). The slides were air-dried for at least 30 min and fixed for 10 mins in 100% methanol (Fisher Scientific, A412SK-4). Methanol fixed cells were stained in May-Grünwald solution (Sigma-Aldrich, MG500-500mL) for 10min and washed 2 times with UltraPure Distilled Water (Fisher Scientific, 10977-023). The cells were then stained for 10 minutes with 10% Giemsa (Sigma-Aldrich, 48900-500mL-F) diluted in UltraPure Distilled Water. The cells were washed 2 times UltraPure Distilled Water and left to air-dry overnight. The slides were mounted with Entellan New (Millipore, 1079610100) after air-drying, and representative pictures were taken using an Axio Lab.A1 microscope (Zeiss) under a N-Achroplan 100x/01.25 objective.

Droplet Digital PCR of sorted cells:

Population 3 cells from day 3 WT or RUNX1-null EBs were sorted into Eppendorf tubes containing FACS buffer, as described above. The FACS isolated cells were centrifuged at 400G for 5 minutes, and the cells were resuspended in 5µL ddPCR 2X supermix (Bio-Rad, catalog# 1863026). Droplet generation was performed on a QX200 ddPCR system (Bio-Rad catalog # 1864001) using cDNA generated from ~500 cells with the One-Step RT-ddPCR Advanced Kit for Probes (Bio-Rad catalog # 1864021) according to the manufacturer's protocol. Reverse transcription incubation was at 42°C and annealing/extension at 60°C. Reactions were partitioned into a median of ~15,800 droplets per well. Plates were read and analyzed with the QuantaSoft software to assess the number of droplets positive for the gene of interest, reference gene, both, or neither. The probers for the assays were purchased from Bio-Rad: HBG2 (AssayID: dHsaCPE5191281), HBE1 (AssayID: dHsaCPE5043916), HBB (AssayID: dHsaCPE5044290), and B2M (AssayID: dMmuCPE5195283).

Statistical analysis

Error bars in graphical data represent means ± SEM. Statistical significance was determined using a Student t test, Mann-Whitney, ordinary one-way ANOVA with Tukey's multiple comparisons test, or Kruskal-Wallis multiple comparisons tests depending on experimental settings and whether the data was normally distributed as determined by a Shapiro-Wilk normality test. $P < 0.05$ was considered statistically significant. Statistical analyses were performed using GraphPad Prism software. The experiments were not randomized, and the investigators were not blinded to allocation during experiments and outcome assessment. All experiments were repeated at least two times to ensure reproducibility of the observations. No statistical methods were used to predetermine sample size.

**Extended Data Figures**

LAZAROV et al., Fig.S1, related to Figure 1.

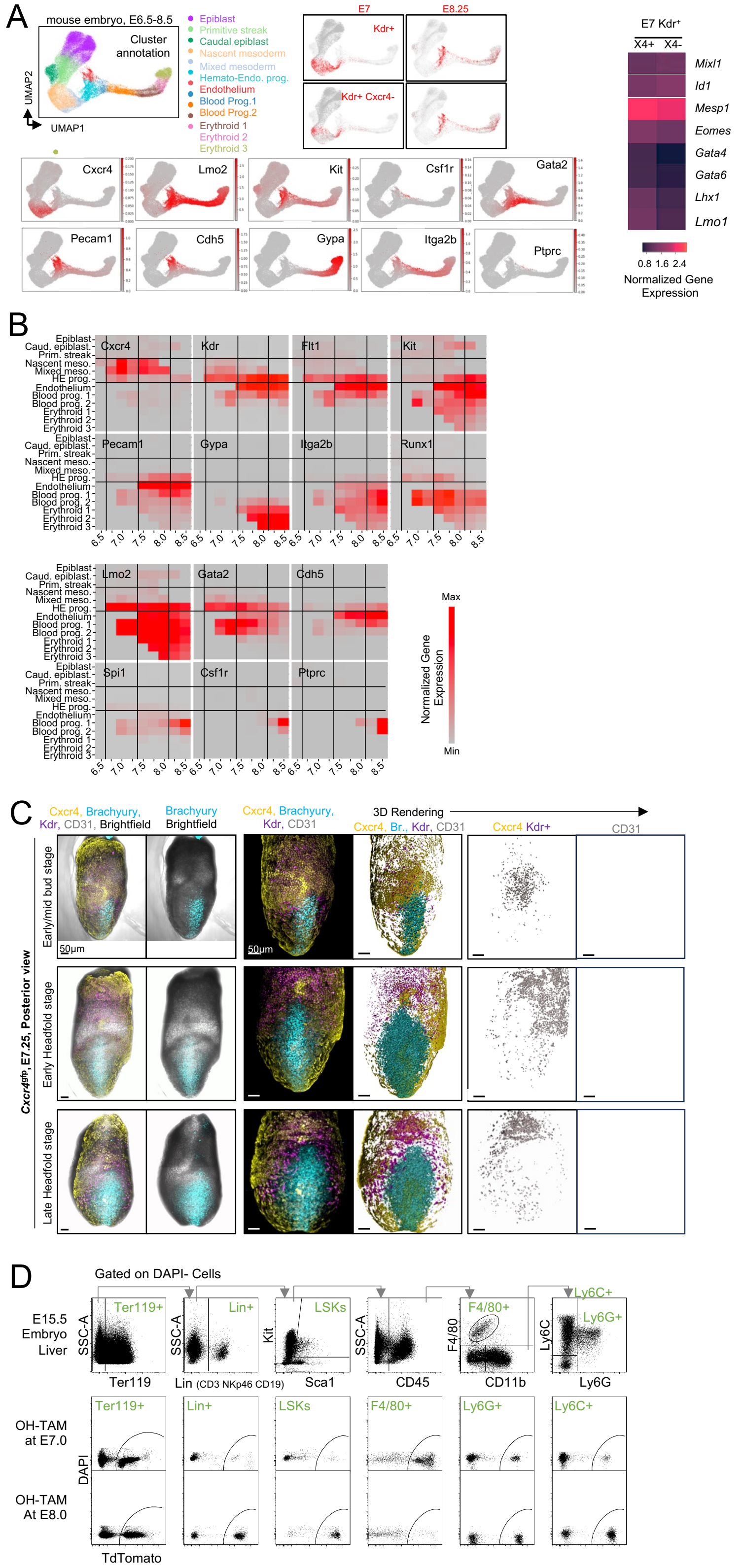


**Extended Data Fig. 1,** related to Figure 1:

**CXCR4 expressing cells in early embryos. A.** Reanalysis of single-cell RNAseq dataset (Pijuan-Sala et. Al, Nature 2019). Left: UMAP scatter plot shows annotated cell types present during mouse gastrulation from E6.5 to E8.5. Endoderm populations have been excluded from the analysis. Grey dots represent all events present from E6.5 to 8.5. Right: Selected DEG between E7 Kdr$^+$Cxcr4$^+$ and E7 Kdr$^+$Cxcr4$^-$ cells. Statistics: Wilcoxon rank sum test with Benjamini-Hochberg correction.
**B.** Reanalysis of single-cell RNAseq dataset (Pijuan-Sala et. Al, Nature 2019). UMAP scatter plots show expression of select mesodermal, endothelial and hematopoietic genes in mouse cells in E6.5 to E8.5 dataset.
**C.** Representative whole mount images of posterior view of Cxcr4$^{GFP}$ embryos at different developmental stages observed for E7.25, and stained with anti-GFP (Yellow), anti-KDR (Purple), anti-Brachyury (Cyan), anti-CD31 (Grey). Analysis of Kdr$^+$Cxcr4$^+$, cells was performed using Imaris software surface-surface statistics (3D Rendering. Scale bars are 50µm. Early/mid bud stage images are from the same embryo shown in figure 1B. **I.**
**D.** Gating strategy for analysis of E15.5 Cxcr4$^{CreERT2}$; R26$^{LSL-tdT}$ embryos pulsed with OH-TAM. Abbreviations: Embryonic day (E); 4-Hydroxytamoxifen (OH-TAM); Uniform manifold approximation and projection (UMAP).

LAZAROV et al., Fig S2, related to Figure 2.

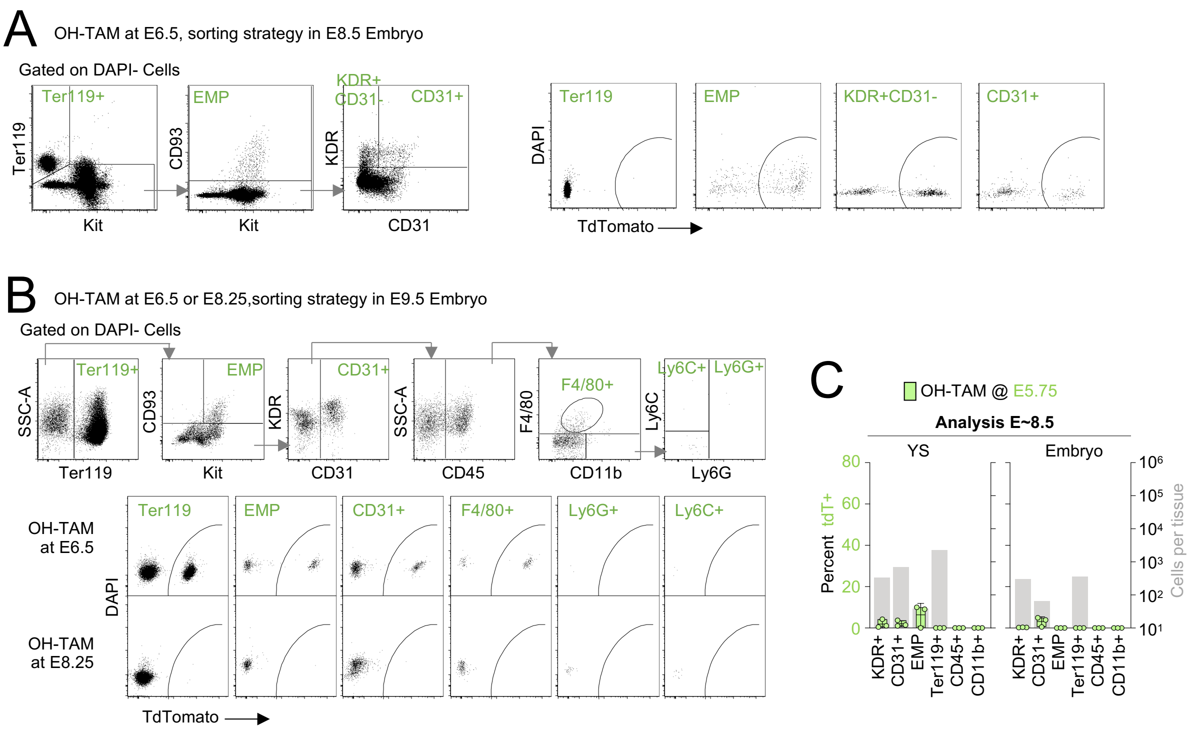


**Extended Data Fig. 2,** related to Figure 2: **Gating strategies for analysis of E8.25 and E9.5 embryos, and differential gene expression analysis of E7 CXCR4$^{+}$KDR$^{+}$ vs CXCR4$^{-}$KDR$^{+}$ cells**. **A, B**. gating of tdT$^{+}$ cells in E8.5 (A) and E9.5 (B) Cxcr4$^{CreERT2}$; R26$^{LSL\text{-}tdT}$ embryo pulsed in vivo with OH-TAM at E6.5 (A,B)or E8.25 (B). **C.** *Cxcr4$^{CreERT2}$; R26$^{LSL\text{-}tdT}$* embryos treated with OH-TAM at E5.75 were analyzed by flow cytometry at E8.5. Green axis: % tdT labeled cells and grey axis: total cell number. N=3 embryos.

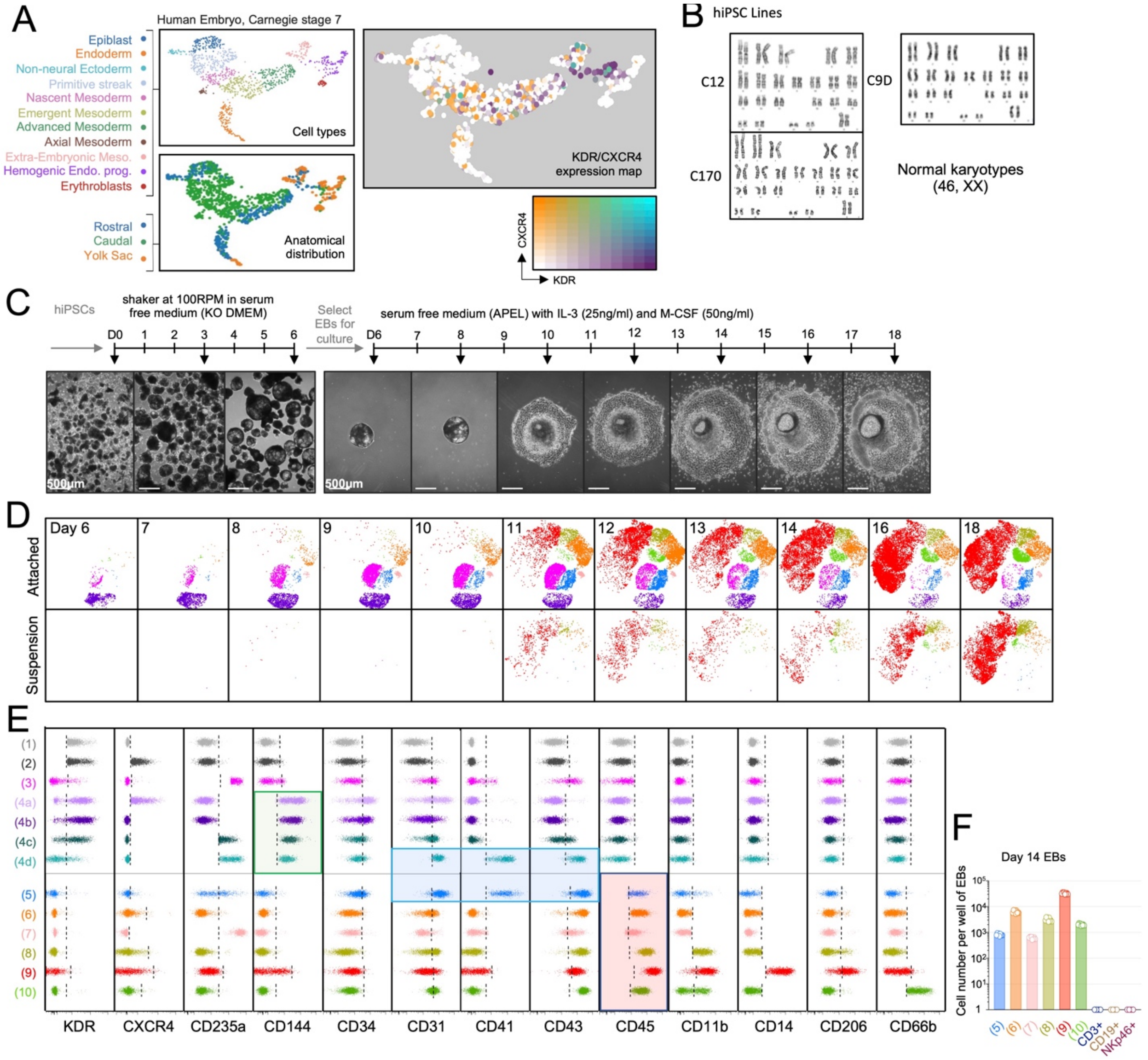


**Extended Data Fig. 3,** related to Figure 3: **Human embryo single-cell RNAseq analysis and characterization of iPSC differentiation model**. **A.** Reanalysis of human scRNAseq data from Human Embryo, Carnegie stage 7 from Tyser et. al., Nature 2021, accessed and analyzed using the tools provided by the authors at: www.human-gastrula.net. Plots show cell types as annotated by the authors, their anatomical distribution, and cells expressing CXCR4 and/or KDR. Abbreviations: Mesoderm (Meso.); Endothelial (Endo); progenitors (Prog.). **B.** Chromosomal analysis of human iPSC lines generated from independent donors used in this study. Normal female (46,XX) karyotype was observed in all DAPI-banded metaphases where n=20 per line. **C.** Schematic of differentiation protocol and representative brightfield microscopy images from indicated days of the differentiation process. hiPSCs aggregates are cultured in KO-DMEM on an orbital shaker from day 0 to day 6 to initiate formation of EBs as indicated in schematic. At day 6 select EBs are transferred in serum free APEL medium on tissue culture plates to allow for their attachment and production of mature hematopoietic cells. **D,E.** Flow cytometry analysis of hiPSC-derived cell populations (see Main Figure 3) from day 6 to day 18 of the differentiation process. (D) tSNE representation of cells whole EBs (Top) and floating cells (Bottom). Colors within tSNE plots correspond to the designated cell populations as shown in figure 3. Number of events in tSNE plots was normalized to 60,000 total live cells (DAPI negative) per timepoint. CXCR4$^{+}$ or

$KDR^+$ cells and events negative for all of the used antibodies are not shown in this tSNE. (E) Expression of indicated proteins in identified cell populations in day 3 and 6 cultures (population 1-4d) and day 15 cultures (population 5-10). Each vertical dotted line represents the maximum fluorescence value of an FMO control for a respective cell population. Panels show representative data where n=3 from 3 independent iPSC lines and 2 independent experiments/line. **F.** Flow cytometry quantification of $CD3^+$, $CD19^+$, or $NKp46^+$ cells in day 14 EBs. n=8 from 2 independent experiments. Abbreviations: human induced pluripotent stem cells (hiPSC); Revolutions Per Minute (RPM); Differentiation Day (D); Embryoid Bodies (EB).

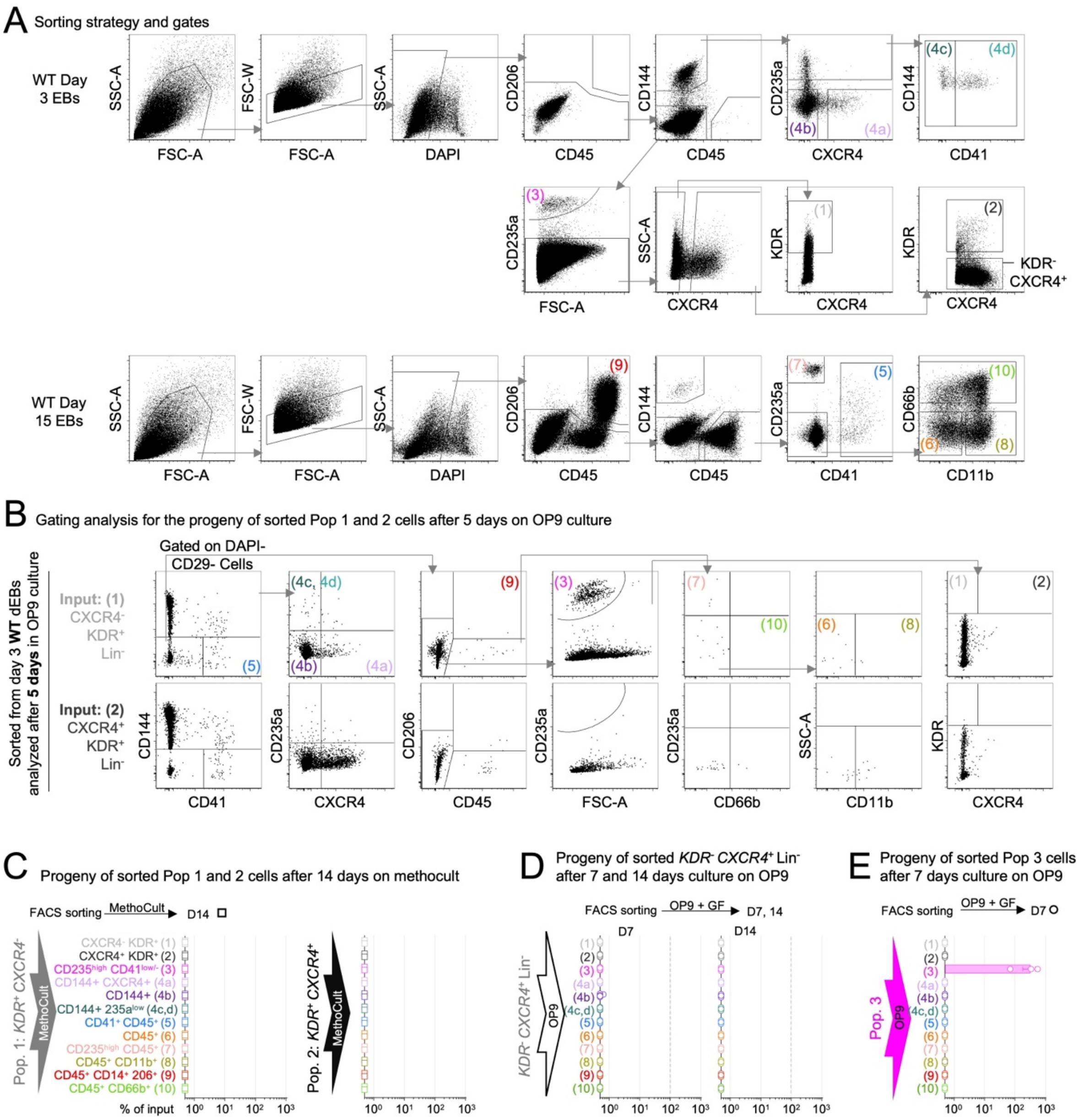

**Extended Data Fig. 4,** related to figure 4. **Gating strategies, and differentiation on OP9 stroma and Methocult**. **A.** Gating strategy for FACS purification of Pops 1 to 10. **B.** Gating strategy for FACS analysis of the progeny of Pop.1 and 2 after 5 days in OP9 cultures. **C.** Progeny of sorted Pop 1 and 2 cells after 14 days on Methocult**:** 100-1000 cells were FACS isolated from day 3 EBs, cultured for 14 days in MethoCult cell potential assay conditions and analyzed by flow cytometry to quantify resulting cell populations as percent of sorted cells input. n = 2 independent experiments. **D.** Progeny of sorted KDR$^-$ CXCR4$^+$ cells (see A), after 7 and 14 days. 100-1000 cells were FACS isolated from day 3 EBs, cultured for 7 and 14 days on OP9 cell differentiation potential assay conditions and analysed as in (C). n = 6 or 5 independent experiments for day 7 and day 14 timepoints respectively. **E.** Progeny of sorted Pop 3. 100-1000 cells were FACS isolated from day 3 EBs, cultured for 7 days on OP9 cell differentiation potential assay conditions and analysed as above. n= 3 independent experiments per cell type.

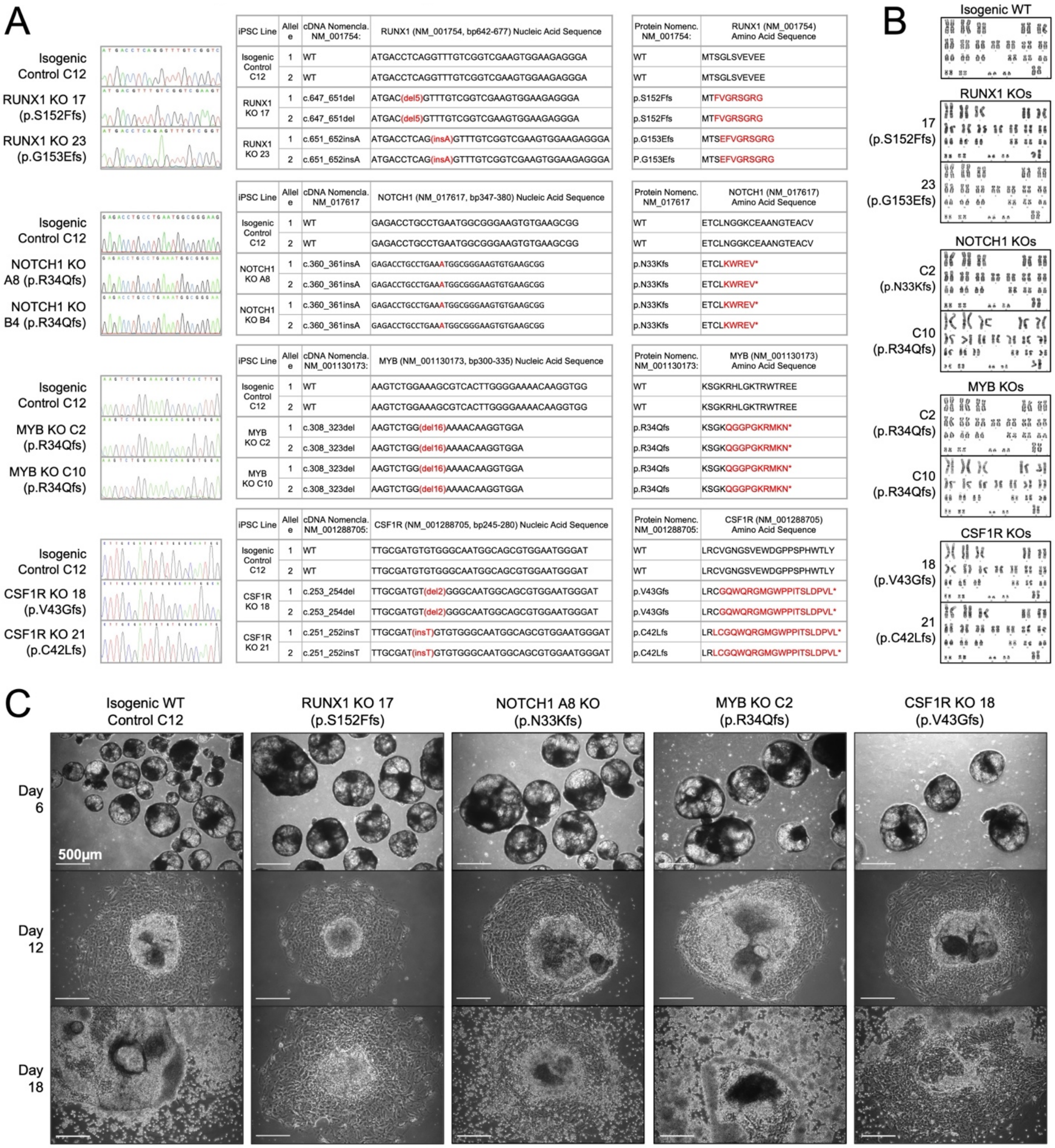


**Extended Data Fig. 5,** related to Figure 4 and 6. **Genetic analysis, karyotype, and differentiation culture brightfield images of CRISPR-CAS9 modified iPSC lines. A.** Sanger sequencing of CRISPR-CAS9 modified iPSC lines and isogenic control. Nucleic acid and predicted amino acid sequences are shown for both alleles based on Sanger sequencing data in CRISPR-CAS9 modified and isogenic iPSC lines. Variations in nucleic acid and amino acid sequences are indicated in red. **B.** Karyotypes of CRISPR-CAS9 modified isogenic iPSC lines. Normal female (46,XX) karyotype was observed in all DAPI-banded metaphases where n ≥ 20 per line. **C.** Representative brightfield images of EBs at day 6, 12, and 18 from isogenic and KO iPSC lines. Abbreviations: induced Pluripotent Stem Cells (iPSC); Nomenclature (Nomenc.); Knock Out (KO).

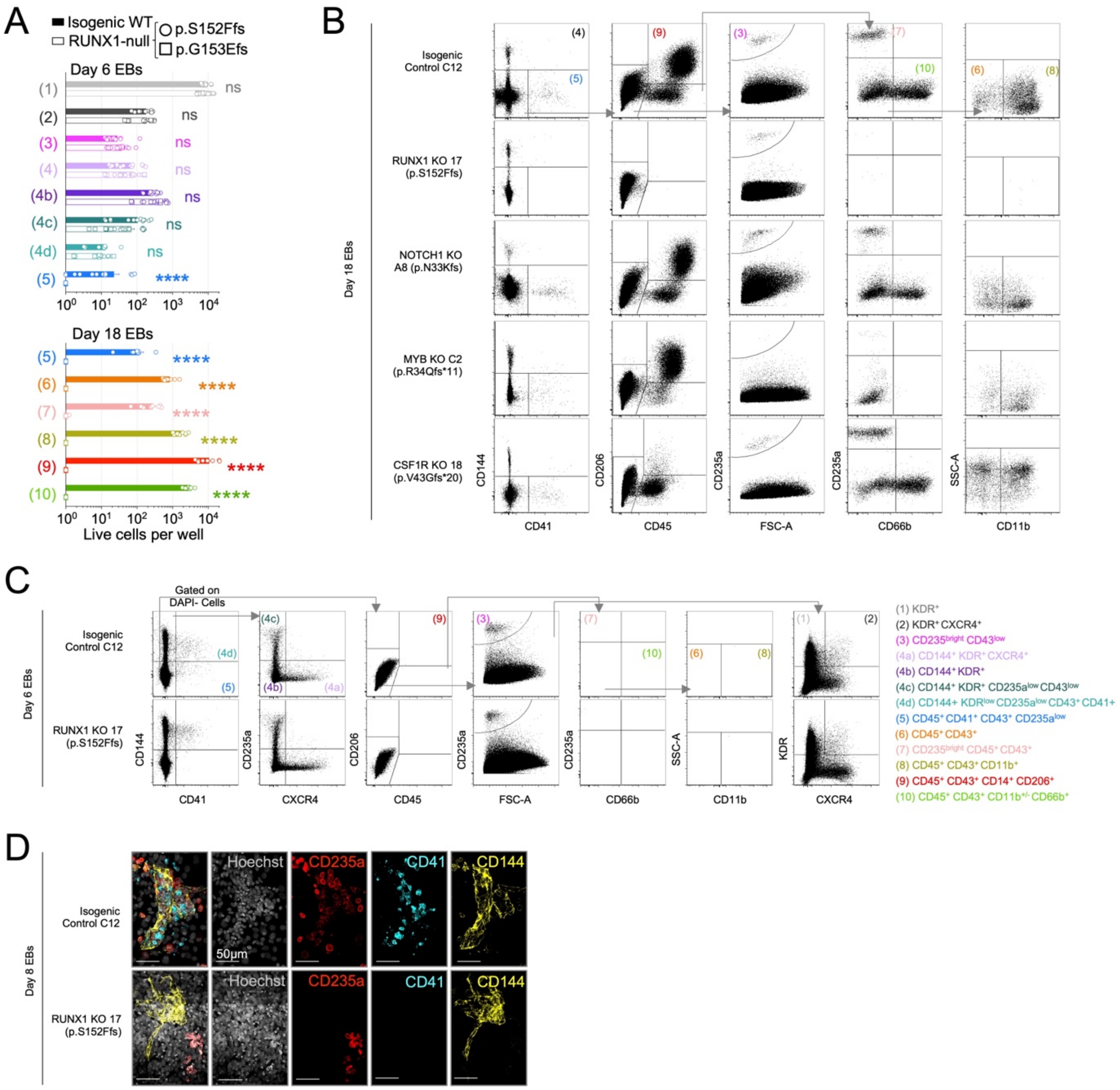


**Extended Data Fig. 6,** related to Figure 4 to 6. **analysis of mutant and isogenic WT cultures.** **A.** Isogenic WT and RUNX1-null (p.S152Ffs and p.G153Efs) iPSC lines were differentiated and analyzed by flow cytometry at day 6 and day 18 to quantify total numbers of the indicated cell populations. n = 8 to 9 from 6 independent experiments. Statistical comparisons were calculated by Mann-Whitney test. not significant (ns); $p \le 0.05$ (*); $p \le 0.01$ (**); $p \le 0.001$ (***); $p < 0.0001$ (****) **B,C.** Two-dimensional flow cytometry examples of representative differentiation cultures in isogenic WT and mutant iPSC lines at day 6 (C) and day 18 (B). Gray arrows indicate gating relationships between plots. Colored numbers indicate the gates for specific cell populations the phenotypes of which are summarized in panel C. **D.** Whole-mount immunofluorescence imaging of a blood island in day 12 Isogenic WT or RUNX1-null (p.S152Ffs) EBs. Representative images from 2 independent experiments.

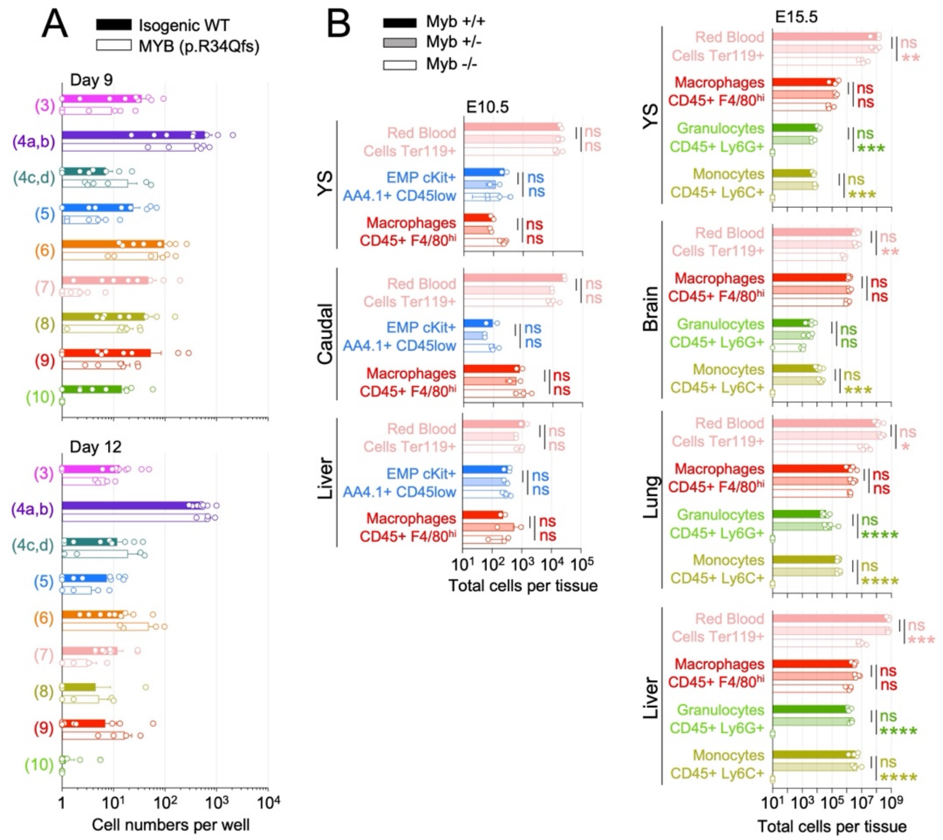


**Extended Data Fig. 7**, related to Figure 4. **Quantification of endothelial and hematopoietic cells in MYB deficient iPSC differentiation cultures and mouse embryos. A.** Isogenic WT and MYB-null (p.R34Qfs) iPSC lines were differentiated and analyzed by flow cytometry at differentiation day 9 and 12 to quantify total numbers of indicated cell populations. n=4 to 9 from 4 independent experiments for each line. **B.** Myb mutant and littermate control mice were analyzed by flow cytometry at E10.5 (Left) or E15.5 (Right) to quantify total numbers of indicated cell populations within the specified tissues. n=2 to 5 mice per timepoint. Statistical differences are calculated with Kruskal-Wallis multiple comparisons test made between groups within each population. Abbreviations: Embryonic day (E); Yolk sac (YS); Erythro-Myeloid Progenitors (EMP); not significant (ns); p≤0.05 (*); p≤0.01 (**); p≤0.001 (***); p<0.0001 (****)

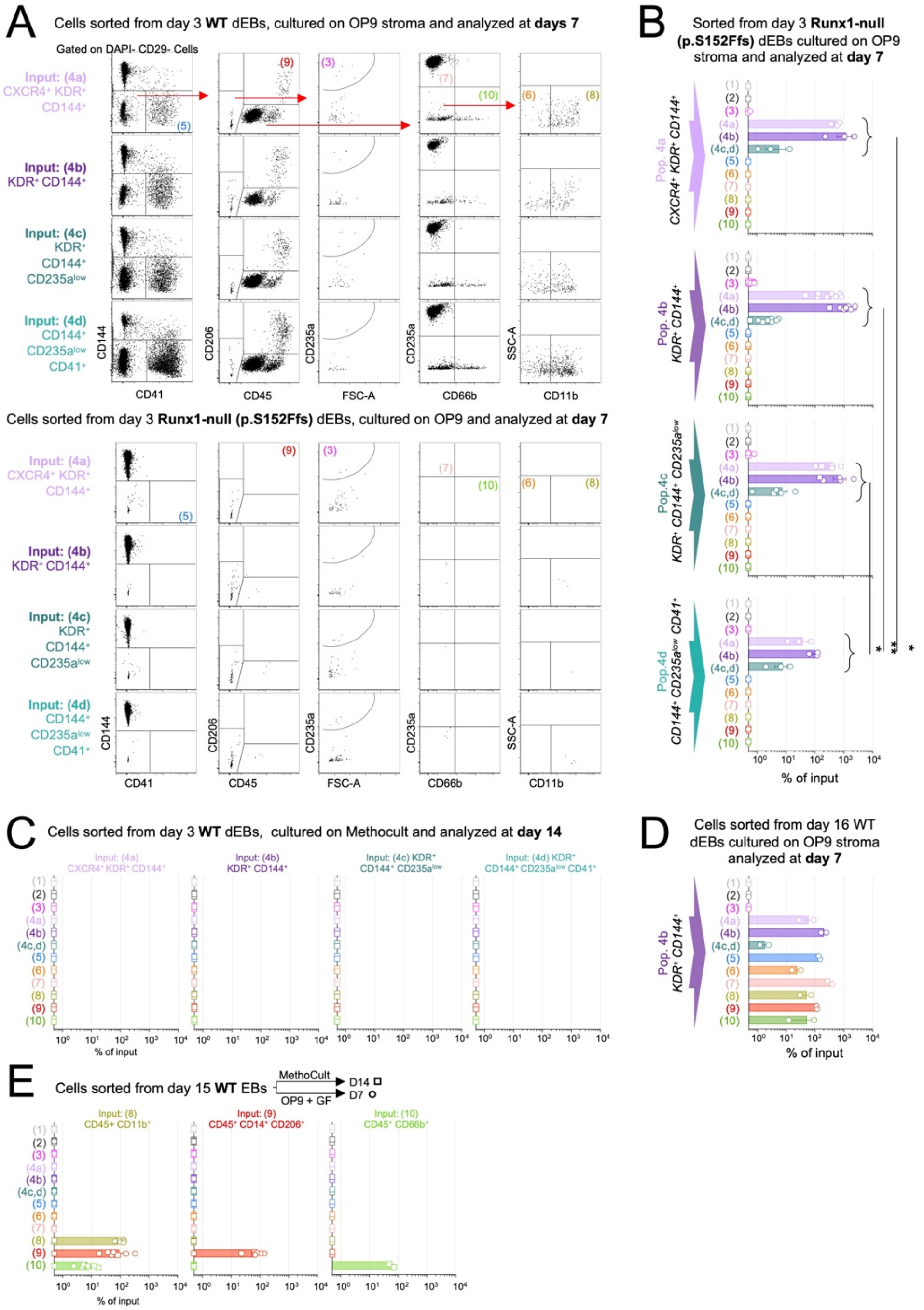


**Extended Data Fig. 8,** related to Figure 4 and 5. **Analysis of Pop. 4 and CD45+ subsets. A.** Flow cytometry plots showing representative examples of day 7 OP9 differentiation potential assay cultures (see methods) with 2,000 FACS isolated cells of population 1, 2, 4a-d from isogenic WT or Runx1-null (p.S152Ffs) day 3 EBs. Arrows indicate gating relationships between plots. Colored numbers indicate the gates for specific cell populations. **B.** 2,000 cells of population 4a, 4b, 4c, or 4d were FACS purified from day 3 Runx1-null (p.S152Ffs) and cultured for 7 days in OP9 cell potential assay conditions (see methods). Whole well cultures were analyzed by flow cytometry to quantify resulting cell populations as a percent of sorted cells input. Statistical comparisons are made between the sum of endothelial cells output (sum of pop 4a-d) from sorted Pop. 4d in comparison to Pop 4a, Pop. 4b or Pop 4c. P values were calculated by Mann-Whitney statistical tests. n=3 to 9 from 3 independent experiments. $p \leq 0.05$ (*); $p \leq 0.01$ (**); $p \leq 0.001$ (***);

p<0.0001 (****). **C.** 100-1000 cells of indicated populations were FACS isolated from day 3 WT EBs and cultured for 14 days in MethoCult cell potential assay conditions (see methods). Whole well cultures were analyzed by flow cytometry to quantify resulting cell populations as percent of sorted cells input. n=3 to 7 from 3 to 6 independent experiments. **D.** OP9 differentiation potential assay for population 4b isolated from day 16 WT EBs. n=2 from 2 independent experiments. **E.** 100-1000 cells of indicated populations were FACS isolated from day 15 WT EBs and cultured for 7 days in OP9 cell potential assay conditions (circles), or 14 days in MethoCult cell potential assay conditions (squares). Whole well cultures were analyzed by flow cytometry to quantify resulting cell populations as percent of sorted cells input. n=3 to 7 from 3 to 6 independent experiments.
